# Mapping the latent CRBN-molecular glue degrader interactome

**DOI:** 10.64898/2025.12.02.691773

**Authors:** Pius Galli, Shuhao Xiao, Yanxiang Meng, Alexander Hanzl, Alexandra M. Bendel, Jacob D. Aguirre, Anna M. Diaz-Rovira, Maximilian R. Stammnitz, Georg Kempf, Lukas Kater, Kenji Shimada, Ye Wei, Andreas Scheck, Dominique Klein, Simone Cavadini, Guillaume Diss, Bruno E. Correia, Nicolas H. Thomä

## Abstract

Molecular glue degraders (MGDs) are a transformative modality in drug discovery. MGDs that work in concert with the E3 ligase CRL4^CRBN^ can degrade a range of substrates through tailored MGDs. To explore CRL4^CRBN^ reprogrammability, we tested whether reported CRBN-MGD substrates are part of a network of latent CRBN interactors, detectable with generic CRBN-MGDs. Leveraging highly parallel interaction measurement (GluePCA) between CRL4^CRBN^ and human zinc-fingers (ZFs), we identified ∼210 ZFs bound to CRBN-pomalidomide, where top binders are already reported as degraded by dedicated MGDs. To map latent CRBN-MGDs interactions proteome-wide, and thus define the immediately accessible CRBN target space, we combined AI-derived protein surface queries (MaSIF-mimicry) with GluePCA. This pipeline identified 6 known and 43 novel CRBN-pomalidomide binders, thereby providing privileged starting points for MGD development. We expect this binding-focused, highly parallel workflow to be readily applicable to other MGD/E3 ligase systems, extending the target landscape of this emerging drug class.

## INTRODUCTION

Molecular glue degrader (MGD) drugs, such as lenalidomide (Revlimid), have revolutionised the treatment of multiple myeloma (MM)^1^, targeting zinc-finger (ZF) transcription factors (TFs) Ikaros and Aiolos for degradation through the CRL4^CRBN^ E3 ubiquitin ligase^2–7^. Lenalidomide derivatives are founding members of the family of MGD drugs, which engage target proteins through their conserved glycine β-hairpin α-turn (also known as G-loop). The Ikaros, Aiolos Cys_2_-His_2_ (C_2_H_2_) ZFs belong to the largest class of human TFs with more than 675 members, all highly related in structure^8,9^. In recent years, a steadily growing number of ZFs, including Helios^10,11^, WIZ^12,13^, ZBTB16^14^, and SALL4^15,16^, has been identified as being degraded by CRL4^CRBN^ through bespoke MGD derivatives. In addition to ZFs, other protein classes and folds, such as GSPT1^6^, VAV1^17^, and kinases (CK1α, etc.)^4,7^, also serve as MGD-dependent substrates of CRL4^CRBN^. The majority of these contain a G-loop motif, present in more than 2,500 proteins in the human genome^18^. Recently, non-G-loop motifs have been reported, further extending the target space^17,19,20^. The wide potential reach of these MGDs is contrasted by their apparent specificity in cellular degradation^21,22^. Hence, a key question for the development of novel therapeutics is which proteins, or folds, are potential CRBN-MGD targets and how specificity can be achieved.

Historically, MGD discovery has been driven by phenotypic screening of large chemical libraries^23^. Recent approaches include degradation reporters^20,24^, mapping CRBN-MGD interactomes by mass spectrometry^17,19,25^, and computational approaches^17,19^. Thus far, there is no integrated, scalable workflow available to comprehensively map the CRBN-MGD target space proteome-wide. Here, we present a highly parallel pipeline to computationally and experimentally identify latent CRBN-MGD interactions, including substrates beyond the canonical G-loop motifs. We utilise a combined workflow of highly parallel interaction measurements (GluePCA) and surface-based computational tools (MaSIF-mimicry) to identify, proteome-wide, which proteins are already measurably engaged with CRBN-MGD through generic, existing MGDs and thereby constitute *bona fide* starting points for the development of specific degraders.

## RESULTS

### GluePCA measures compound-dependent ZF binding to CRBN

We hypothesise that many proteins found to be degraded through MGD derivatives are already CRBN-engaged through generic MGDs, yet not sufficiently to drive degradation. Such latent proximity inducers are prime candidates for degraders, as evident from a number of studies on CRBN as well as other E3 ligases correlating cellular degradation with biochemical binding properties, reporting that a ∼2-3 fold increase in target binding to CRBN-MGDs suffices to convert CRBN-MGD-bound proteins into robustly degraded ones^24,26^. Hence, knowing which proteins latently interact with CRBN-MGD E3 ligase complexes, even if not yet degraded, will inform CRBN target discovery.

To identify CRBN-MGD-bound proteins on a large scale requires overcoming the limitations of traditional biochemical approaches, allowing us to profile thousands of protein interactions. We chose a highly parallel protein complementation assay (PCA)^27^, where a methotrexate-resistant murine dihydrofolate reductase (DHFR) is split into two fragments (“DH” & “FR”), which are fused to bait and prey proteins. The binding between the proteins reconstitutes DHFR activity, allowing yeast growth to serve as a proxy for interaction (**Fig. 1a** and see **Methods**)^28^. We set out to test whether DH-bait and FR-prey complementation can also be induced by MGDs (GluePCA^29^). We first chose CRL4^CRBN^ and the ZF family of TFs, where the FR-fragment was fused N-terminally to the ZF, and the DH-fragment to CRBN (**Fig. 1a**). As a positive control, we tested an Ikaros construct (ZF2, residues 141-174), the minimal domain required for CRBN binding in the presence of pomalidomide^24^, which showed sustained growth, indicating binding between Ikaros ZF2 and CRBN in the presence of pomalidomide. In contrast, no binding was observed in the absence of pomalidomide, confirming that GluePCA is a suitable assay for the CRBN system (**Fig. 1b**), albeit higher compound concentration is required in yeast than is the case in mammalian cells (**Extended Data Fig. 1c)**. To quantify and compare growth between single-clone GluePCAs, we computed the area under each curve (AUC)^30^.

**Figure 1:**
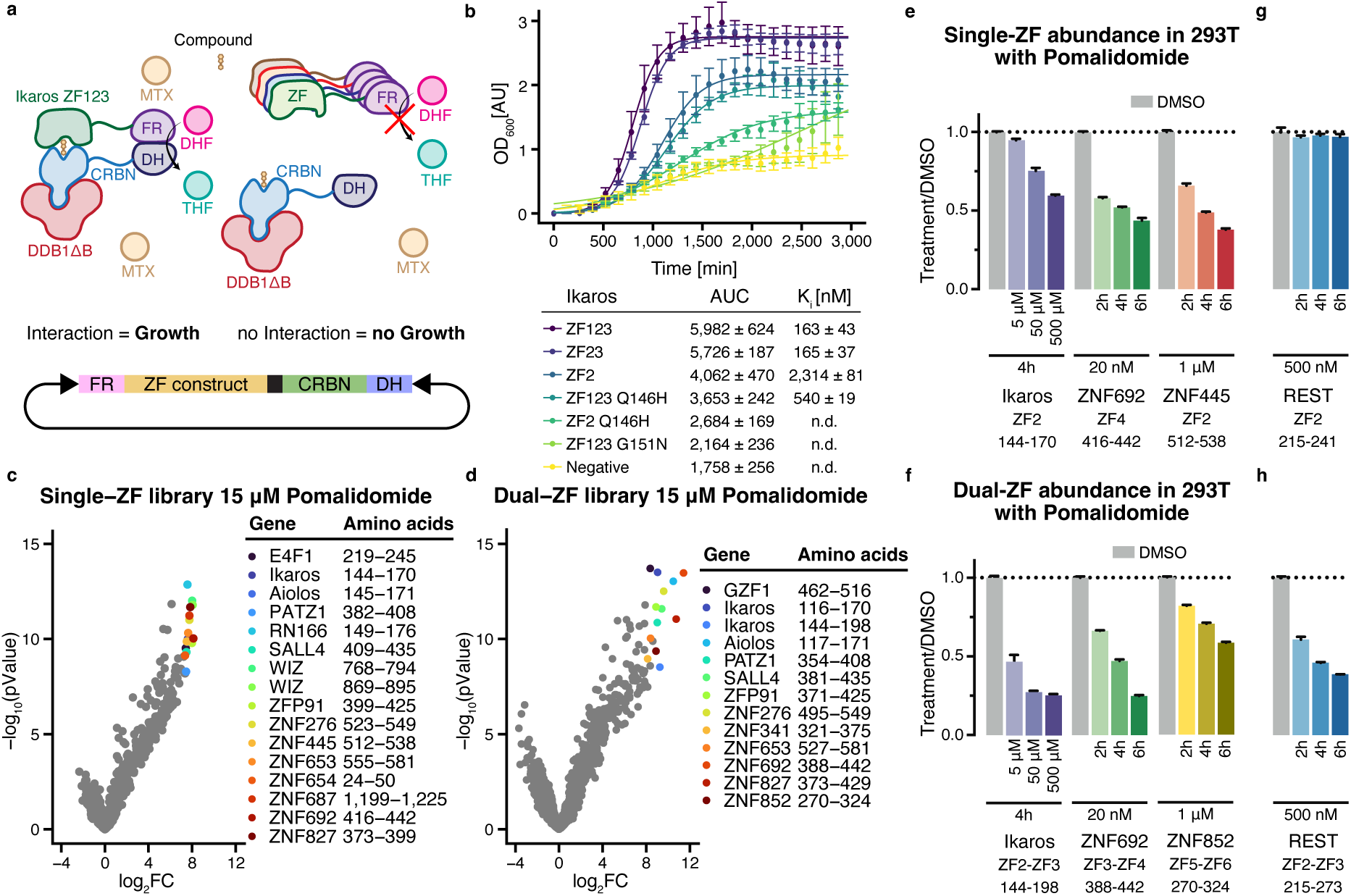
Proteome-wide GluePCA screen identifies over 100 binding and 2 novel degrading ZF constructs in the presence of pomalidomide. **(a)** Schematic of the GluePCA and the vector used. FR-/DH- both are N-terminally fused to the CRBN and ZF construct with a terminator in the middle. THF: Tetrahydrofolate, DHF: Dihydrofolate. **(b)** Single-clone GluePCA in yeast, constitutively expressing DDB1ΔB, with FR-Ikaros ZF constructs and DH-CRBN, grown for 2,888 min in 100 µL selective media (SC-Ura/Ade/Met) containing pomalidomide and Methotrexate (MTX). OD_600_ was measured every 15 min in a plate reader and normalized with a blank (Biological replicates = 3, each point indicates the mean of the experimental replicates, error bars indicate the standard deviation (SD)) **(c)** & **(d)** Log_2_FC of sequencing read counts (pomalidomide/DMSO) and corresponding pValue (calculated with limma) for the Single ZF (c) and dual ZF (d) library with raw read count > 10 in all three replicates (Biological replicates = 3) **(e), (f), (g)** & **(h)** 293T cells expressing the indicated ZF-GFP-IRES-mCherry construct were treated for 2, 4 and/or 6h with DMSO or pomalidomide at the indicated dose. Cells were analysed by flow cytometry to quantify the GFP over mCherry ratio, which was normalised to DMSO for each treatment (Biological replicates = 3, bar heights indicate mean of experimental replicates, error bars indicate one standard deviation)

Next, we assessed the dynamic range of the assay with different lengths of Ikaros constructs, spanning ZF1 to ZF3 **(Extended Data Fig. 1b)**, and Ikaros mutants. Ikaros ZF2-ZF3 (residues 141-195) and ZF1-ZF2-ZF3 (residues 117-195) showed fast growth analogous to stronger binding, compared to ZF2 alone. An IKZF1 Q146H mutant exhibits weaker growth by 2 to 3-fold, while the G151N mutation showed no growth above background **(Fig. 1b)**. These growth rates mirror previous *in vitro* biophysical IC_50_ measurements for CRBN complexes (IC_50_ in probe displacement: ZF1-ZF2-ZF3: ∼460 nM, ZF2-ZF3: ∼470 nM, ZF2: ∼7,3540 nM)^24^. GluePCA thus provides a rapid and sensitive readout of MGD-induced CRBN binding, matching the dynamic range of biophysical assays while eliminating the need for protein purification.

Encouraged by these results, we expanded GluePCA to pooled high-throughput screens. We screened a library of 8,363 individual C_2_H_2_ ZFs across 1,407 uniquely expressed genes and quantified binding propensities using next-generation sequencing **(Extended Data Fig. 1b)**. This led to the identification of numerous ZF targets previously reported to be degraded by pomalidomide, including Ikaros/Aiolos, RNF166, WIZ, ZFP91, ZNF276, ZNF653, ZNF692, and ZNF827^3,5,13,24,31^ **(Fig. 1c)**. Comparing our data to published single ZF degradation screens, we identified 9 out of 11 previously reported ZFs^24^. Among the top binders, we also identified SALL4, a validated pomalidomide target^15,16^, which eluded previous single ZF degradation screens^24^. ZNF445^20^ was recently identified in a dual ZF degradation screen; notably, we also observed degradation of its single ZF in cells **(Fig. 1e)**. Out of the top 16 hits, the remaining two ZFs (PATZ1, ZNF687)^31,34^ have been reported to be degraded in cells as full-length (FL) proteins. We also observed Helios, for which biochemical recruitment with pomalidomide had been previously observed^6,24^; yet, degradation required compound modification^10,11^. To further experimentally validate GluePCA hits, we posited that true hits would show dose-dependency. Using a log_2_FC threshold > 2 and pValue < 0.01 at 15 µM, all 212 hits showed dose-dependent binding **(Extended Data Fig. 1d)**. These findings experimentally confirm previous predictions that more than 100 ZFs are compatible with CRBN-pomalidomide binding, with top binders being subjected to degradation^24^.

The majority of ZFs in proteins in the human proteome are found as multidomain tandem repeats of single ZF domains^35^ **(Extended Data Fig. 1b)**. While the ZF contacting the CRBN-MGD interface is sufficient for binding and degradation, adjacent ZFs can strengthen the interaction and trigger robust degradation^20,24^. The contribution of these flanking ZFs remains unclear. We refer to ZFs directly engaging the CRBN-MGD interface as main ZFs (mZFs), and preceding and subsequent ZFs as accessory ZFs (aZFs).

To dissect how multiple ZFs engage CRBN-MGD complexes, we designed a library containing all tandem C_2_H_2_ ZFs in the human proteome with a 6-7 residue spacing **(Extended Data Fig. 1b)**, yielding 5,293 dual ZFs across 713 unique genes. To experimentally define true binders, we repeated GluePCA at a lower pomalidomide concentration. 285 dual ZF binder constructs with log_2_FC > 2 showed dose-dependent binding **(Extended Data Fig. 1e)**. Overall, the top dual ZF hits aligned with the single ZF screen, including ZNF692, ZNF827, Ikaros/Aiolos, ZNF276, SALL4, PATZ1, ZFP91, and ZNF653, yet the rank-order changed **(Extended Data Fig. 1f)**. We also identified previously unreported pomalidomide-recruited ZFs: ZNF852 and ZNF341, a TF regulating STAT3^36^. In cellular ZF degradation reporters, both showed robust degradation with pomalidomide **(Fig. 1f and Extended Data Fig. 1c)**. We further observed REST, a transcriptional repressor implicated in multiple cancers^37–40^, bound CRBN-pomalidomide as a single ZF. While REST ZF2 alone was not degraded, the dual ZF2-ZF3 construct was degraded in the presence of pomalidomide **(Fig. 1g, h)**, underscoring the importance of aZFs in driving degradation.

Having established GluePCA as a highly parallel binding assay to identify CRBN-pomalidomide targets across thousands of ZFs, we confirm the existence of a continuum of ∼200 dose-dependent CRBN binders. The top hits were either previously reported as degraded by pomalidomide or showed robust degradation only with optimised compounds (i.e., FIZ1 with Compound 1^41^, ZBTB11 with WJ-01-306^42^, Helios with NVP-DKY709^11^ and ALV2^10^). The findings for REST further illustrate that the ZF degron that governs binding to CRBN-MGD complexes is more complex than a single ZF and that additional ZF interactions facilitate degradation.

### CRBN-pomalidomide recognises a three-consecutive ZF degron

To dissect how multiple ZFs engage the CRBN-MGD complex, we generated a deep mutational scanning (DMS) library of 89,918 single-point mutations from 124 dual ZF binders. We subjected this library to GluePCA. For most constructs, one ZF was dominant, and mutating this ZF showed the most deleterious effect on binding **(Fig. 2a and Extended Data Fig. 2a, b and 3a, b)**. This dominant ZF typically also scored better than its neighbouring partner ZF in single ZF GluePCA **(Extended Data Fig. 3c)** and was hence assigned mZF. DMS analysis revealed that while some ZFs were largely unaffected by N- or C-terminal aZF mutations, others showed enhanced or reduced binding **(Fig. 2a and Extended Data Fig. 3a)**. The data suggested individual mutations can affect both ZF-fold integrity and CRBN interaction. Averaging the aZF mutation profiles did not reveal a conserved, easily identifiable binding patch.

**Figure 2:**
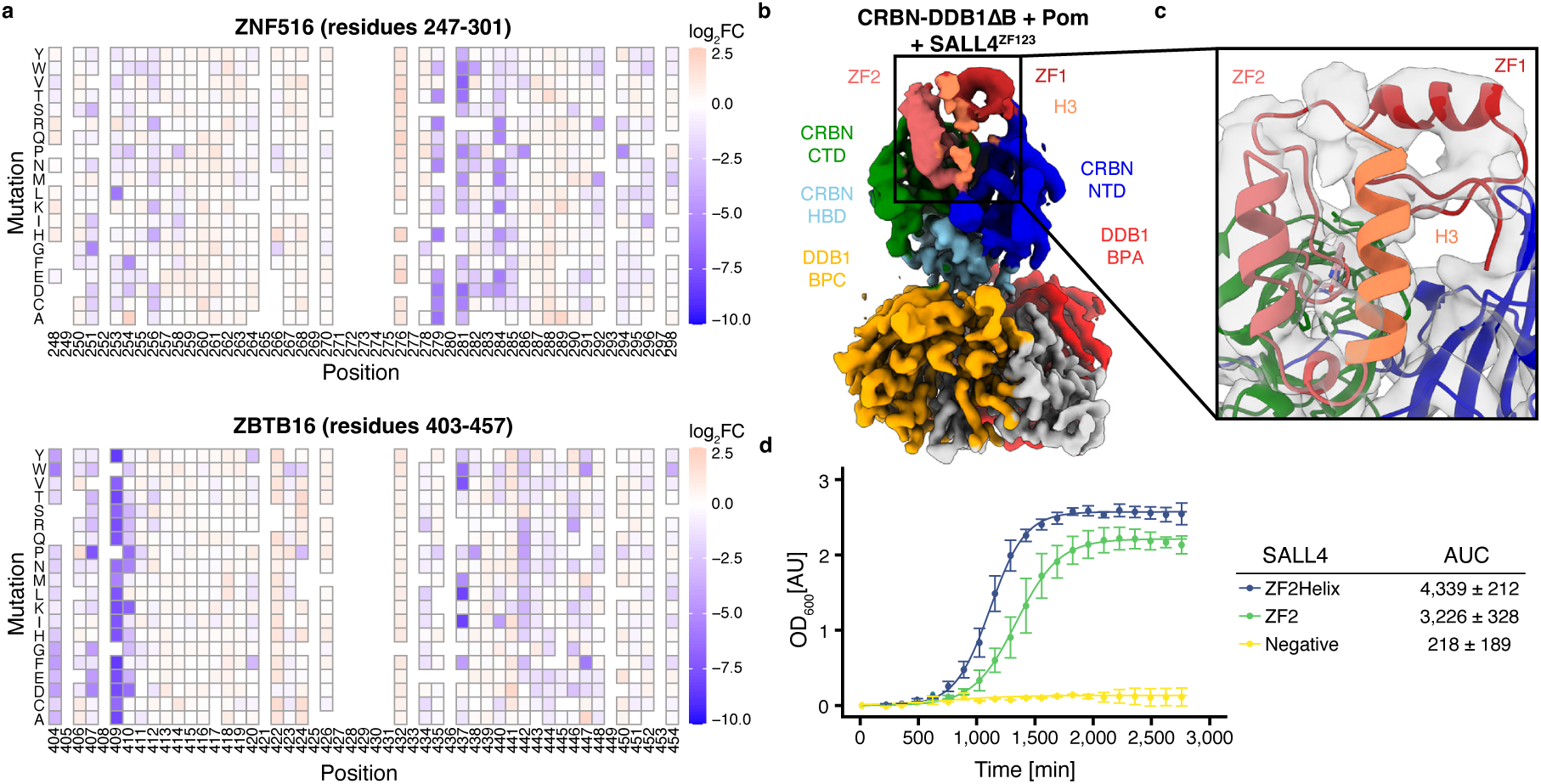
The N- and C-terminal accessory ZF enhances selective binding. (a) Example heatmaps of the deep mutational screen for ZNF653 and ZNF516 (DMS) of the DMS library in GluePCA. (b) Cryo-EM structure of SALL4^ZF123^ bound to DDB1ΔB∼CRBN and pomalidomide. CTD, carboxy-terminal domain; HBD, helical bundle domain; NTD, amino-terminal domain; BPA & BPC, β-propeller A and C, respectively. (c) Close-up of SALL4 ZF1, ZF2, and the helix fitted into our cryo-EM density (grey). (d) Single clone GluePCA in yeast with SALL4 ZF2Helix and ZF2 construct. ZF2 contains residues PQVKA (residues 433-436) preceding the helix, which were shown to benefit binding. (Biological replicates = 3, each point indicates the mean of the experimental replicates, error bars indicate the SD)

**Figure 3:**
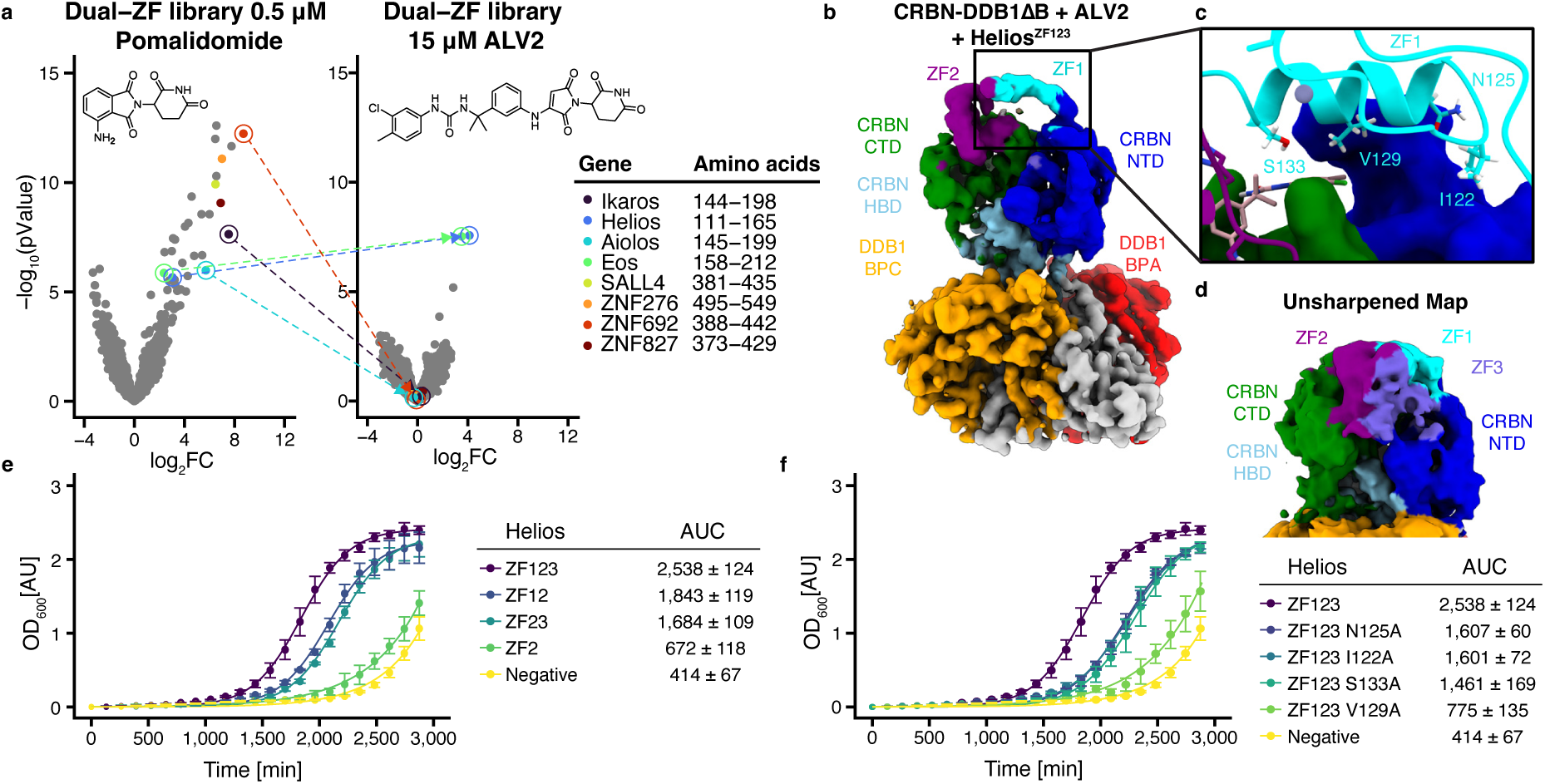
Compounds can alter specificity through direct interaction with the accessory ZF. (a) Log_2_FC of sequencing read counts (pomalidomide or ALV2/DMSO) and corresponding pValue (calculated with limma) for the dual ZF library with raw read count > 10 in all three replicates (Biological replicates = 3). (b) Cryo-EM structure of Helios^ZF123^ bound to DDB1ΔB∼CRBN and ALV2. (c) Close-up of Helios ZF1. (d) Unsharpened density of the Helios cryo-EM structure with ZF3 visible in violet (e) Single-clone GluePCA in yeast with Helios ZF123, ZF12, ZF23 and ZF2 construct. (Biological replicates = 3, each point indicates the mean of the experimental replicates, error bars indicate the SD) (f) Single-clone GluePCA in yeast with Helios ZF123, N125A, S133A, I122A and V129A mutants. (Biological replicates = 3, each point indicates the mean of the experimental replicates, error bars indicate the SD)

To identify potential binding sites for the aZF on CRBN binding, we reconstituted two dual ZFs: SALL4 aZF1-mZF2 and Ikaros mZF2-aZF3. Cryo-EM structures of these ZFs bound to DDB1^ΔB^, CRBN, and either SALL4^ZF1-ZF2-ZF3^ or Ikaros^ZF2-ZF3^ in the presence of pomalidomide had an overall resolution of 3.5 Å **(Extended Data Fig. 4a-e and 5a-e and Table S1)**. The SALL4 structure found aZF1-mZF2, in a position consistent with the previously reported X-ray structure^43–45^ **(Fig. 2b)**. Interestingly, we observed additional density C-terminal of mZF2 resembling a sole α-helix absent from the crystallographic construct, not previously observed (yet predicted in AlphaFold2 (AF2)^46^) **(Fig. 2c)**. Single-clone GluePCA assays showed that the presence of the helix strengthened binding ∼1.4-fold **(Fig. 2d)**. The Ikaros^ZF2-ZF3^ map also revealed density beside mZF2 which we assigned to aZF3 **(Extended Data Fig. 3d)**. While we see the SALL4 N-terminal aZF1 interacting with CRBN NTD residues F102, I152, H353 and F391, the Ikaros C-terminal aZF3 interacts with CRBN NTD residues Q86 and H102 (**Extended Data Fig. 3e, f)**. Based on the SALL4 and Ikaros structure, we conclude that the aZF N- or C-terminal of mZF can comprise up to three consecutive ZFs (aZF-mZF-aZF). In addition, our DMS screen suggests that the aZFs (N- or C-terminal) can either facilitate or weaken binding, as would be expected for neomorphic CRBN-MGD interactions^47^ **(Extended Fig. 3a)**. While the mZF binding mode is relatively conserved, the aZF contributes through multiple and apparently non-conserved binding modes in line with a neomorphic, evolutionary non-conserved, binding mode.

### The molecular basis of specificity for CRBN-based degraders

In this study, we see several ZFs bound with pomalidomide, a first-generation thalidomide derivative. However, degradation for FIZ1, ZBTB11 and Helios is only evident upon further compound optimisation. This poses two questions: how does the compound favour the selected target, and, assuming MGDs are largely specific, how does the compound prevent the remainder of the ∼200 ZFs from being degraded?

To examine the basis of specificity, we performed GluePCA experiments with Helios-specific ALV2 degrader using the dual ZF library, in conjunction with solving the cryo-EM structure of CRBN bound to Helios in the presence of ALV2. In GluePCA, the most enriched members were Helios and the related Eos (IKZF4), confirming specificity, with a more substantial enrichment of Helios than in experiments with pomalidomide. The most unexpected effect of ALV2, however, was the significant drop in enrichment for nearly all other ZFs in the library, including Ikaros, ZNF692, and SALL4 **(Fig. 3a)**. ALV2 selectivity hence involves stabilising the target and destabilising the remainder of the ZF target space.

To define the structural basis for selectivity, we solved the cryo-EM structure of DDB1^ΔB^, CRBN, and Helios^ZF1-mZF2-ZF3^ with ALV2 at 3.6 Å resolution **(Fig. 3b and Extended Data Fig. 6a-e and Table S1)**. The structure shows Helios mZF2 and aZF1 in a conformation previously reported for ALV1^10,20^, with aZF1 resembling SALL4 aZF1 as well as weak density for aZF3 **(Fig. 3c, d)**. We then used single-clone GluePCA to dissect the binding contributions of both aZFs. The presence of both aZF1 and aZF3 exhibited stronger binding compared to mZF2 alone **(Fig. 3e)**. While ALV2 affinity is achieved by accommodating Helios H141 at the mZF as previously reported, the model further revealed Helios aZF1 residues I122, N125, and V129 interacting with CRBN residues H103, P104, F150, I152, while Helios S133 interacts with ALV2 **(Fig. 3c)**. Accordingly, in GluePCA we found that mutation of I122A, N125A, S133A reduced binding, while V129A ablates the interaction **(Fig. 3f)**, explaining stabilisation beyond mZF2 through the aZFs.

A comparison between the Helios/ALV2 **(Fig. 3b)** and SALL4/pomalidomide **(Fig. 2b)** structures provided insight into selectivity and specificity beyond the mZF. SALL4 did not show ALV2 binding in our screen **(Fig. 3a)**, consistent with predicted aZF clashes with the compound **(Extended Data Fig. 6f)**. Superimposing other ZF-CRBN structures, such as ZNF692, similarly not found engaged with ALV2 in our assays, also finds the compound blocking ZNF692 aZF1 binding, with leucine (L413) replacing the critical Helios serine (S133) that engages ALV2. Given the ALV2 selectivity for Helios in the dual ZF library, combined with the structures, suggests ALV2 stabilisation of the mZF together with disfavouring binding of other ZFs with the help of the accessory ZFs.

### Proteome-wide prediction of CRBN targets using surface mimicry

Using a highly parallel screen, we identified more than 200 detectable, compound-dependent CRBN-pomalidomide binders in a single fold class. Most of these ZF binders identified here, along with other known CRBN substrates, share a β-hairpin G-loop motif but lack a clear sequence signature beyond a conserved glycine **(Extended Data Fig. 7a)**. The promiscuous CRBN-MGD binding observed suggests additional, largely uncharted CRBN interactions may exist across the proteome even with generic MGDs such as pomalidomide. We first developed a computational pipeline to narrow the search scope and, through protein engineering, enable high-parallel experimental GluePCA validation at low cost. This approach was rooted in two central computational resources: *(i)* a structural database of 75,684 domains from AF2-predicted structures for structure-guided searches, and *(ii)* a surface-based structural alignment tool for rapid comparison of protein surfaces to find similar patches which may mediate similar interactions. The latter tool was based on the MaSIF^48^ framework, referred to as MaSIF-mimicry (**Fig. 4a and** see **Methods)**. State-of-the-art co-folding methods, such as AlphaFold3 (AF3), are computationally expensive to scale at the proteome level and struggle to predict novel MGD-mediated ternary complexes^49^, partly due to the absence of evolutionary constraints in neomorphic interfaces. We therefore asked whether surface-based representations could bypass sequence and structural constraints (e.g., β-hairpin G-loop) to identify surface features resembling known binding interfaces, even in structurally and evolutionarily distant proteins.

**Figure 4:**
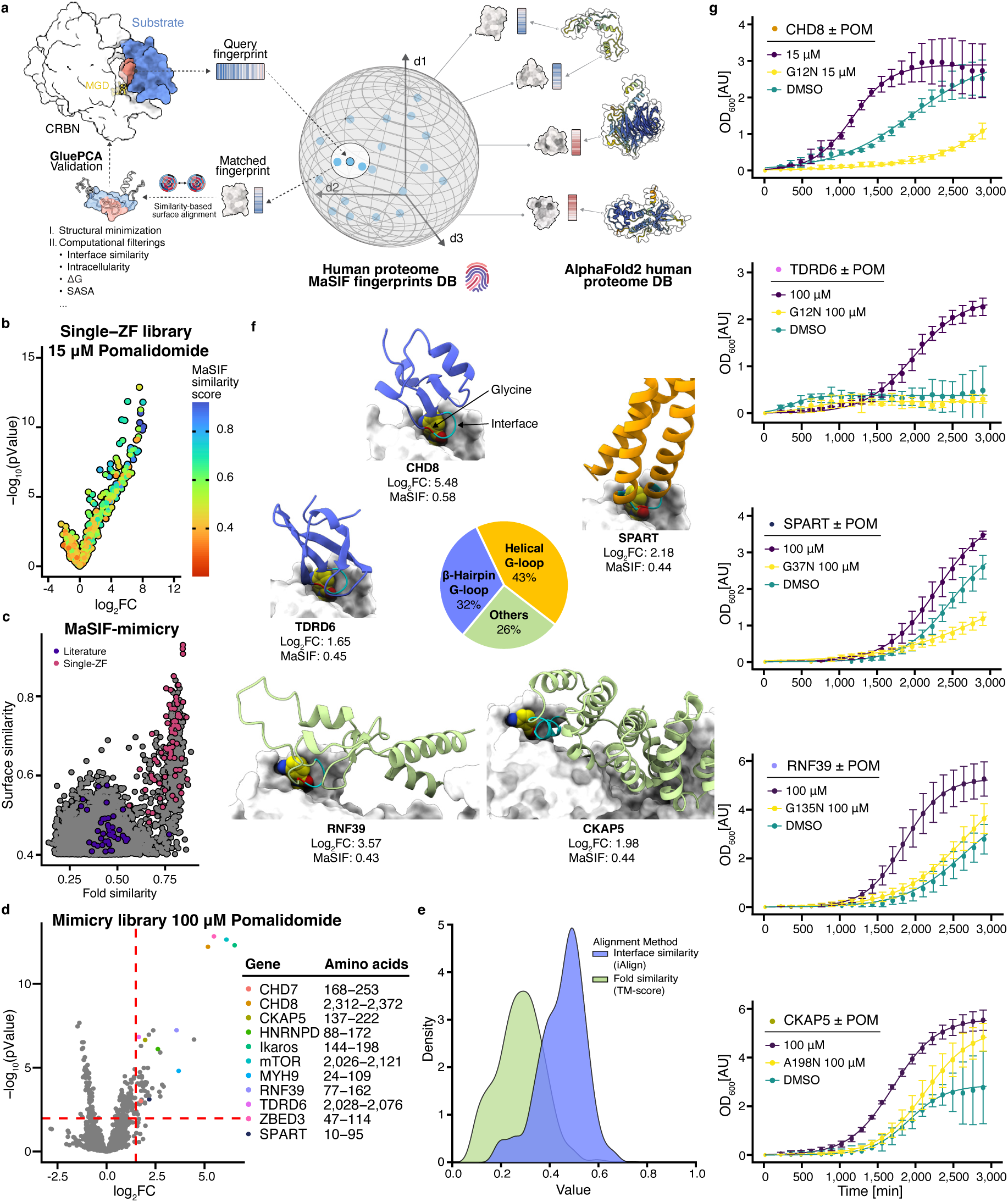
An *in-silico* surface mimicry search can predict novel CRBN-MGD binders and expand the CRBN target space. **(a)** Schematic of the MaSIF-mimicry search pipeline. **(b)** Volcano plot from the Single ZF library colored with interface similarity to that of ZNF692 ZF4 (red = low; blue = high) **(c)** Proteome-wide MaSIF-mimicry screen with Ikaros ZF2. The x-axis indicates fold similarity (TM-score >0.5 is considered as strong structural similarity), while the y-axis indicates interface similarity. Red: known hits from the Single ZF library; violet: known hits from literature, including Baek et al.^18^ and Petzold et al.^16^. **(d)** Log_2_FC (pomalidomide/DMSO) for variants with raw counts > 10 across all replicates (Biological replicates = 3; thresholds: Log_2_FC > 1.5, p < 0.01 **(e)** Score distribution of all-against-all similarity measurements of predicted interfaces and the fold between all 50 binders identified from the *bonsai* library, including Ikaros ZF2. **(f)** Predicted binding modes for CHD8, SPART, and CKAP5 (cyan = Interface; orange = helical; blue = β-hairpin) **(g)** Single-clone GluePCA validation. AUC (WT, Mut, DMSO): CHD8 5,128 ±812, 790 ±40, 3,221 ±418; CKAP5 6,915 ±281, 4,310 ±837, 3,112 ±884; SPART 2,801 ±424, 1,149 ±226, 1,784 ±290; RNF39 5,845 ±1.039, 2,717 ±353, 2.031 ±624; TDRD6 2,384 ±303, 622 ±157, 799 ±752 (Biological replicates = 3; points = replicate means; error bars = SD).

We benchmarked MaSIF-mimicry using the single ZF library to test if it recovers GluePCA hits by querying the surface patches surrounding G423 of ZNF692 ZF4 (PDB ID: 6H0G) against all ZF domains in the human proteome. Strikingly, most known CRBN-pomalidomide ZF substrates (i.e., Ikaros, SALL4, and WIZ) share strong similarity (MaSIF similarity score > 0.4) with the ZNF692 interface despite low sequence homology, indicating ZF substrates share common binding surfaces despite diverging sequence. Thus, MaSIF-mimicry can retrospectively uncover novel ZF substrates proteome-wide using AF2 structures in the apo-state **(Fig. 4b and Extended Data Fig. 7b)**. Other homology-based metrics (e.g., sequence similarity, interface pseudo-sequence similarity, ESM2 embedding distances, or structural similarity) showed far weaker discrimination **(Extended Data Fig. 7b)**.

We next expanded the search space to the entire human proteome to test whether MaSIF-mimicry can identify evolutionarily distant proteins outside of the ZF protein family as potential drug targets for CRBN-mediated target protein degradation. Bound interfaces of known neo-substrates (CK1α, Ikaros, ZNF692, and GSPT1) were queried against the AF2-predicted human proteome database. Notably, in each case, a substantial number of domains were found to have similar sites to these G-loop interfaces. Thus, CRBN’s apparent promiscuity can be attributed to the widespread presence of a similar surface pattern across the proteome, while also suggesting the existence of a largely unexplored latent CRBN interactome. Among these computational hits, several previously reported substrates were recovered, including CHD7 and PPIL4, which were found by mass spectrometry^19^, as well as NEK7 and MNAT1, through proximity-ligation^17^ **(Fig. 4c)**. We also retrieved VAV1^17^ with GSPT1 as query **(Extended Data Fig. 7c)**, and despite the divergent secondary structure at the interface, the known mTOR CRBN-binder is successfully identified in ZNF692 mimicry searches **(Extended Data Fig. 7c)**. Importantly, the predicted binding mode generated by MaSIF-mimicry for VAV1, mTOR, and PPIL4 are consistent with experimentally resolved structures^17,19^ **(Extended Data Fig. 7c)**. This illustrates that by performing rapid, proteome-wide MaSIF-mimicry searches we can identify shared interface patterns from structurally distant proteins proteome-wide. While these proteins are only found degraded by specialised compounds, we computationally identify them already as binders through mimicry of substrates bound to CRBN-pomalidomide.

To prospectively identify new CRBN-MGD binders, we screened a library comprising 1,959 top-ranked non-ZF candidates (based on MaSIF similarity score) from the initial 7,480 protein domains with surface sites similar to the Ikaros ZF2 interface (PDB: 6H0F) identified by MaSIF-mimicry. GluePCA achieves its highest quantitative accuracy when candidate proteins with similar geometries are present at comparable lengths and abundances, as this facilitates interaction between the DH-and FR-fragments with similar expression levels and geometry, enabling direct comparison among binders. This, along with the ensuing cost of FL gene synthesis, led us to design a library where we reduced the length of the candidate binder to the minimal sub-domain that includes the degron required for CRBN interaction. Due to the constraints from gene synthesis, larger proteins were truncated *in silico* to sub-domains containing 86 residues around the interface (we refer to this as a *bonsai* library). These truncated sub-domains, along with proteins smaller than 86 residues, were filtered based on structural and bioinformatic criteria to exclude transmembrane segments, intrinsically disordered regions, and potentially unstable truncations (see **Methods and Extended Data Fig. 7d, e)**. After this in silico workflow, the 1,959 highest-scoring protein domains were retained and screened by GluePCA, including Ikaros as a positive control.

From this Ikaros-mimicry bonsai library subjected to GluePCA, we recovered several known neo-substrates, including mTOR^17^, ZBED3^19,51^, CHD7^19^, HNRNPD^17^, and MYH9^19^ **(Fig. 4d)**. We also identified NEK2, a homologue of NEK7^17^, predicted to interact with CRBN-pomalidomide in a binding mode similar to NEK7 (PDB: 9H59; RMSD 3.5 Å), again where NEK7 was previously identified only through degradation by dedicated MGDs (**Extended Data Fig. 7c**). Beyond these, 43 novel protein domains showed compound-dependent binding (log_2_FC > 1.5, pValue < 0.01), spanning diverse protein folds with predicted similar surface signatures at the interface **(Fig. 4e and Table S2)**. Notably, based on MaSIF-mimicry, several of the hits are not predicted to bind CRBN-MGD *via* the β-hairpin G-loop interface. Some utilise helical G-loops (e.g., SPART), or even bypass the crucial glycine entirely (e.g., CKAP5) **(Fig. 4f)**. Using single-clone GluePCA, we validated two β-hairpin G-loop binders as full domain (FD) (CHD8, a transcriptional repressor linked to autism^53^, and TDRD6) and one helical G-loop binder (SPART FD), in all cases G-to-N mutants disrupted binding **(Fig. 4f**). Both CHD8 and SPART bind in the absence of pomalidomide. Pomalidomide markedly enhances CHD8 binding, whereas SPART shows only a subtle dose-dependent increase. We also validated RNF39 (FD), predicted to contain a G-loop yet classified as neither β-hairpin nor helical G-loop, by introducing a G-to-N mutation. To validate CKAP5, a microtubule-associated protein promoting cell proliferation and cisplatin resistance in esophageal squamous cell carcinoma^52^, we examined the predicted binding interface, which features an alanine in place of the glycine in canonical G-loops. We reasoned that an A-to-N mutation, analogous to the G-to-N mutation, should weaken binding. As expected, the A-to-N mutation weakens binding, validating CKAP5 as a novel non-G-loop CRBN target **(Fig. 4g).** Many GluePCA hits found in this study represent hard-to-drug proteins^54^ **(Table S2)**; in addition, they cover a diverse set of domains, like PDZ, RRM, and WW, across multiple protein families (e.g., kinases, plakins, helicases) and biological processes, such as transcriptional regulation, cell differentiation, and beyond, illustrating the reach of the CRBN-MGD system **(Table S2 and Extended Data Fig. 7f)**.

Our results demonstrate that combining an *in silico* MaSIF*-*mimicry screen with a high-throughput experimental validation backend provides an actionable workflow for expanding the latent CRBN-MGD interactome. This enables the identification of novel CRBN-MGD *bona fide* targets, which, given pre-existing affinity, are prime targets for protein degrader development.

## DISCUSSION

The CRL4^CRBN^ E3 ligase has emerged as an unexpectedly versatile degrader system. This raises three central questions: *(i)* which additional human proteins can be targeted given the appropriate MGD; *(ii)* how these diverse substrates can be engaged; and *(iii)* how selectivity is achieved through tailored MGDs.

To address these questions, we established a scalable pipeline combining computational screening (MaSIF-mimicry) with experimental validation (GluePCA) to uncover latent CRBN-MGD interactions. Using the ZF family as a first test case, the largest human TF family^8,9^, typically considered undruggable, we identified more than 210 ZFs as CRBN-MGD binders in dose-dependent manner, tripling the number of known putative targets. Furthermore, we revealed that most of the newly identified neosubstrates are surface mimics of other previously known ones; based on this, MaSIF-mimicry searches with the Ikaros interaction motif revealed more than 7,000 potential intracellular binders, which were then filtered and truncated into stable domains for high-throughput GluePCA screening. From 1,959 top-ranked hits, 43 novel protein constructs and 6 known substrates were identified as dose-dependent CRBN-pomalidomide binders. Beyond the canonical β-hairpin G-loop and the recently discovered helical G-loop, we identified 9 constructs predicted to engage CRBN in a G-loop-independent manner. We consider the *bonsai* approach a coarse pre-screen used to validate MaSIF-mimicry predictions when the intact domain or FL proteins cannot be screened. Together, these findings highlight how GluePCA, combined with surface-centric computational approaches, can effectively identify potentially druggable targets across distant protein families at scale, independent of sequence or structural homology. Importantly, as new CRBN-MGD-target structures and compounds emerge, MaSIF-mimicry can be rapidly expanded to new substrates and new protein-interface space, which GluePCA can further validate.

Mechanistic analyses revealed that ZFs engage CRBN via degrons spanning up to three tandem-repeat ZFs, sometimes incorporating accessory elements such as the SALL4 helix. A central mZF thereby interacts directly with the compound, while flanking aZFs form weaker CRBN interactions. These aZF interactions, consistent with their neomorphic character, can either stabilise or destabilise binding, often determining degradation. Exceptions such as WIZ^12^, which contains ZF domains widely spread apart acting independently, highlight diverse degron architectures beyond the three-ZF and two-ZF-helix motifs described here. Growing out the compound towards the ZF enables direct mZF and aZF interactions that provide specificity and affinity (e.g., ALV2, **Fig. 3b**)^17^. Future MGD designs will have the possibility to exploit accessory-domain/MGD interactions, thus enhancing potency and selectivity.

Recent proteomic approaches, including IP-MS^18^ and TurboID^16,48^, have expanded the CRBN substrate landscape. GluePCA, by virtue of being an overexpression format, assures that candidate proteins are present in the target cell, including complete mutational control of the interactions for validation and perturbation. Computational methods based on MaSIF have also been applied in focused studies, such as the identification of VAV1 as a non-G-loop CRBN neosubstrate^16^. Our strategy differs in that we employ a surface-centric, proteome-wide screening framework to identify, at scale, proteins capable of engaging CRBN-MGD complexes in binding modes analogous to known substrates. Although the same MGD scaffolds may not degrade many such proteins—for example, we identified mTOR from the Ikaros-mimicry bonsai library as a binding protein for CRBN-pomalidomide while its degradation is only observed in the presence of a different MGD^17^—they represent promising candidates for future compound optimisation. While mimicry searches based on substrates binding to CRBN-pomalidomide complexes have already revealed a rich binding landscape, as more chemically diverse MGDs binding more structurally diverse substrates become available, these can then be easily integrated to further extend the space of possible CRBN-MGD interactors.

In summary, we present an integrated workflow that couples MaSIF-mimicry with GluePCA to define the interactome of a given target. Applied here to CRBN-pomalidomide, this approach provides a generalizable strategy to explore and expand the degradation potential of other MGD systems. The identical workflow is fully portable and can be applied to other E3 ligase MGD complexes for which target structures are available. We anticipate that such hybrid proteome-wide profiling approaches will be crucial in expanding the target landscape of MGDs and contribute to enhancing the therapeutic potential of this transformative drug modality.

## Acknowledgment

We thank M. Schütz-Stoffregen and F. Bello for laboratory management, organisation, and assistance with manuscript editing, and Ben Lehner and C. Soneson for discussion. We are grateful to all former and current members of the Thomä lab and Correia lab. This work was supported by funding from the European Research Council (ERC) under the European Union’s H2020 research program (NucEM, no. 884331), the Swiss National Science Foundation (SNF 310030_301206 and 310030_214852), from Krebsforschung (KFS-5933-08-2023), The Mark Foundation for Cancer Research and the Novartis Research Foundation to N.H.T.; the SNSF Swiss Postdoctoral Fellowship (TMPFP3_224735) to A.H; the EMBO Postdoctoral Fellowship (ALTF 142-2024) to Y.M. We thank the Spanish Ministry of Science and Innovation for funding A.M.D.-R.’s predoctoral fellowship (FPU/03921). We also thank EMBO for the Scientific Exchange Grant (n° 10784) to support A.M.D.-R.’s stay at EPFL.

## Data availability

All DNA sequencing data have been deposited in the Gene Expression Omnibus (GEO) and will be accessible with accession number <u>GSE298800</u>. The electron density reconstructions and final models have been deposited in the Electron Microscopy Data Bank (EMDB) and the PDB. They will be accessible with the following IDs: SALL4: 9SQ4, EMD-55106; Ikaros: 9SQ5, EMD-55107; and Helios: 9SQ6, EMD-55108.

## Code availability

The code for the sequencing analysis will be accessible at https://github.com/LabThoma/Latent_CRBN-MGD_interactome. The surface mimicry code will be accessible at https://github.com/LPDI-EPFL/masif_seed in a new branch termed *mimicry*.

## Contributions

P.G., S.X., B.E.C. and N.H.T. conceived the study. D.K. constructed the yeast strain used in this study with the help of P.G. and K.S. P.G. performed the GluePCA experiments, including library preparation and data processing and analysis, with the help of A.M.B. and G.D. P.G. performed the single growth GluePCA experiment, including sample preparation and data analysis, with the assistance of A.M.B., J.A. and K.S. A.H. performed the cellular degradation assays, including data analysis. P.G. purified recombinant DDB1ΔB∼CRBN, SALL4^ZF12Helix^, Ikaros^ZF23^, and Helios^ZF123^ with the help of J.A. P.G. performed the cryo-EM experiments, including sample preparation and data processing, with the help of S.C., L.K., G.K. and J.A. for model building and data acquisition. S.X., A.S., and B.E.C. conceived the surface mimicry pipeline. In collaboration with A.M.D.-R. and Y.W., S.X. developed the algorithm for surface mimicry. S.X. conducted the surface mimicry screen and analysis under the supervision of B.E.C. Y.M. conceived and conducted the filter pipeline for the bonsai library. N.H.T. supervised the structural and biochemical work, as well as the GluePCA assays. P.G., S.X., B.E.C. and N.H.T. wrote the manuscript with input from all authors.

## Corresponding author

Correspondence to Nicolas H. Thomä.

## Methods

### Yeast Strain

All experiments were performed in BY4742 (MAT α his3Δ1 leu2Δ0 lys2Δ0 ura3Δ0) with an ADH promoter-driven DDB1ΔB, integrated at the HPT1 locus with a Hygromycin B resistance marker (hereafter referred to as BY4742 DDB1ΔB).

To integrate ADH-DDB1ΔB at the HPT1 locus, we constructed a plasmid named pAG-ADH-DDB1ΔB with the Gibson Assembly Master Mix (New England Biolabs, Ipswich, MA), with a 3x molar excess of the ADH and DDB1ΔB insert and 50 ng of vector. Next, the integration cassette was PCR amplified with Q5 polymerase (New England Biolabs, Ipswich, MA), column purified with QIAquick PCR Purification Kit (QIAGEN, Hilden, Germany), transformed into BY4742 yeast cells, and plated on YPAD plates containing 150 μg/mL Hygromycin B. Finally, integration of the DDB1ΔB gene was confirmed by colony PCR and Sanger sequencing.

### Media and buffer recipes

- YPAD (1 L): 50 g/L YPAD Broth ready to use (Formedium #CCM1010), filter-sterilised.
- SC-Ura (1 L): 6.7 g/L Yeast Nitrogen Base without amino acids (Formedium #CYN0410), 20 g/L D(+)-Glucose (Formedium #GLU04), 0.77 g/L CSM Single Drop-out -Ura (Formedium #DCS0061), filter-sterilised (Millipore Express ®PLUS 0.22 μm PES, Merck, Darmstadt, Germany).
- SC-Ura/Ade/Met (1 L): 6.7 g/L Yeast Nitrogen Base without amino acids (Formedium #CYN0410), 20 g/L D(+)-Glucose (Formedium #GLU04), 0.77 g/L CSM Triple Drop-out -Ade -Met -Ura (Formedium #DCS0901), filter-sterilised (Millipore Express ®PLUS 0.22 μm PES, Merck, Darmstadt, Germany).
- SORB: 1 M Sorbitol, 100 mM LiOAc, 10 mM Tris-HCL pH 8.0, 1 mM EDTA pH 8.0, filter-sterilised (Millipore Express ®PLUS 0.22 μm PES, Merck, Darmstadt, Germany).
- Plate mixture: 40% PEG3350, 100 mM LiOAc, 10 mM Tris-HCl pH 8.0, 1 mM EDTA pH 8.0, filter-sterilised (Millipore Express ®PLUS 0.22 μm PES, Merck, Darmstadt, Germany).
- Recovery medium: YPAD medium with 0.5 M sorbitol, filter-sterilised (Millipore Express ®PLUS 0.22 μm PES, Merck, Darmstadt, Germany).
- Competition medium: SC-Ura/Ade/Met medium with 200 μg/mL methotrexate (Chemie Brunschwig AG), 2 % DMSO.

### Small-scale yeast transformation

Yeast cells from the BY4742 DDB1ΔB strain were streaked on YPAD plates from a glycerol stock and incubated for two days at 30°C. A single colony was used to inoculate 10 mL of liquid YPAD and was grown at 30°C and 200 rpm overnight until the culture reached saturation. In the morning, OD_600_ was measured, and 10 mL of YPAD was inoculated at OD_600_ = 0.3 and grown at 30°C and 200 rpm for approximately 4 h until an OD_600_ of 1.0 to 1.2 was reached. Cells were collected by 5 min centrifugation at 3,000 g and washed once with 10 mL of sterile ddH_2_O and once with 10 mL of TEL (100 mM LiOAc, 10 mM Tris-HCl, pH 7.5, 1 mM EDTA, filter-sterilised). Then, the sample was resuspended in 100 μL TEL. To this, 10 μL of 10 mg/mL pre-boiled salmon sperm DNA (Agilent Technologies, Santa Clara, CA) was added, and the mixture was thoroughly mixed before adding 100 ng of plasmid. Next, 300 μL of 45% PEG4000 in TEL was added, thoroughly mixed, and the mixture was incubated for 30 min at room temperature (RT) without agitation. The cells were heat-shocked for 10 min at 42°C, spun down for 2 s at 21,300 g, and the supernatant was removed by aspiration. The cells were then washed twice with 1 mL of sterile ddH_2_O. Finally, the cells were resuspended in 100 μL SC-Ura, of which 50 μL was plated on SC-Ura plates and incubated for two days at 30°C.

### Single-clone GluePCA

The experiment was performed in three replicates with three blanks, which contained Competition medium without cells. For each replicate, a single colony of BY4742 DDB1ΔB cells transformed with a plasmid was used to inoculate 10 mL of SC-Ura/Ade/Met and was grown at 30°C and 200 rpm overnight until saturation, typically for 12 to 14 h. Next, the OD_600_ was measured and used to inoculate 100 μL of Competition medium ± compound in a 96-well TPP tissue culture test plate (96F, flat bottom). Incubation and data collection were performed with the lid on using a Tecan Infinite 200 Pro plate reader at 29.9°C and 200 rpm (orbital, 1.5 mm amplitude) for 200 cycles (14 min/cycle) with a settling time of 200 ms.

### Single-clone GluePCA data processing

During analysis, the measurements were first transformed into OD_600_ values by subtracting the mean of the blanks and then multiplied by 10 to get the actual OD_600_. Next, the first measurement was disregarded, and measurements 2 to 6 were averaged. Then, the mean and standard deviation (SD) were calculated over the three replicates. The growth was fitted with an R script based on the Growthcurver package^30^, using the logistic equation common in ecology and evolution^55,56^, which gave the number of cells N_t_ at time t.

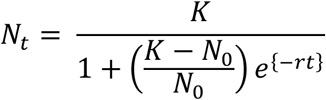

Where N_0_ was the beginning of the growth curve, K the carrying capacity or maximum possible population size, and r the growth rate. Growthcurver was also used to calculate the area under the curve (AUC) for each replicate; finally, the mean and SD of the AUC were calculated.

### Library curation

To curate a list for the Single and Dual ZF libraries, we used PROSITE^57^ (http://prosite.expasy.org/) to scan the proteome for single and dual C_2_H_2_ ZF motifs. The patterns used were: X-X-X-C-x(2,4)-C-x(12)-[CH]-x(3,5)-[CH]-X-X-X (Single ZF) and X-X-X-C-x(2,4)-C-x(12)-[CH]-x(3,5)-[CH]-X-X-X-x(0,1)-X-X-X-C-x(2,4)-C-x(12)-[CH]-x(3,5)-[CH]-X-X-X (Dual ZF). For the Single ZF library, a total of 8,363 ZFs were found in 1,407 genes. The Dual ZF library consisted of a total of 5,293 ZFs in 713 genes. These amino acid sequences were then reverse-translated and codon-optimised with DNA Chisel^58^. Finally, the restriction enzyme sites NheI and HindIII were added together with a common primer site for amplification, and, using DNA Chisel, any introduced NheI and HindIII sites within the coding sequence were removed.

### Library synthesis and cloning

To prepare the backbone for all libraries, we used plasmid pDL5, which was digested with BamHI-HF (New England Biolabs, Ipswich, MA) and SpeI-HF (New England Biolabs, Ipswich, MA), dephosphorylated with Quick CIP alkaline phosphatase (New England Biolabs, Ipswich, MA), gel-purified with the QIAquick Gel Extraction Kit (QIAGEN, Hilden, Germany), and CRBN was then cloned in and fused to the DH fragment. This intermediate plasmid was then digested with HindIII-HF (New England Biolabs, Ipswich, MA) and NheI-HF (New England Biolabs, Ipswich, MA), dephosphorylated with Quick CIP alkaline phosphatase (New England Biolabs, Ipswich, MA), and gel-purified with the QIAquick Gel Extraction Kit (QIAGEN, Hilden, Germany).

All libraries were synthesised by TWIST Bioscience (San Francisco, CA) as oligo pools and PCR-amplified (12 cycles) with KAPA HiFi Kits (Roche). Then, the pools were digested with HindIII-HF (New England Biolabs, Ipswich, MA) and NheI-HF (New England Biolabs, Ipswich, MA) and ligated with NEB T4 ligase (New England Biolabs, Ipswich, MA) into 100 ng of backbone at a 3:1 molar excess of library over backbone. The 20 μL reaction was incubated in a PCR thermocycler with the temperature-cycle ligation (TCL) programme, which consisted of 4 cycles of 10 min at 22 °C and 10 min at 16 °C, followed by 49 cycles of 30 sec each at 4 °C, 27 °C, and 13 °C, then 10 cycles of 30 sec at 4 °C, 1 h at 16 °C, 10 min at 22 °C, and 10 min at 16 °C, another 49 cycles of 30 sec each at 4 °C, 27 °C, and 13 °C, followed finally by 30 sec at 4 °C, 1 hour at 16 °C, 1 h at 22 °C, and 3 h at 18 °C. After TCL, the reaction was dialysed against ddH_2_O using nitrocellulose membranes with 0.025 μm pores (Merck Millipore, Darmstadt, Germany) to reduce salt concentration and concentrated down to 5-10 μL in a SpeedVac. The ligation product was then transformed into NEB 10-beta electrocompetent *E. coli* (New England Biolabs, Ipswich, MA). To increase the yield, three transformations were performed per ligation. For each transformation, 1.5 μL of ligation product was mixed with 25 μL of competent cells, then transferred into a pre-chilled 0.1 cm Gene Pulser Cuvette (Bio-Rad, Hercules, CA). The mixture was electroporated using a Gene Pulser Xcell (Bio-Rad, Hercules, CA) with the exponential protocol at 2.0 kV, 200 Ω, and 25 μF. Next, the cells were immediately resuspended in 488 μL of prewarmed (37°C) SOC medium and the cuvette was rinsed with 488 μL of the same medium. Each transformation was incubated at 37°C for 30 min at 250 rpm. Next, to estimate transformation efficiency, 1 μL of each transformation was diluted 1/1000 and plated onto LB + 200 μg/mL ampicillin with 10 μL and 100 μL of the dilution. The remaining cells were poured onto four LB + 200 μg/mL ampicillin plates. All plates were incubated for 16 h at 37°C. After incubation, the colonies of the dilutions were counted. For each library, an efficiency of > 50 transformants per variant was achieved. The cells were harvested by washing the colonies off the plates with ddH_2_O using a plastic spatula and washed twice with ddH_2_O. Finally, the plasmids were purified with the QIAGEN Plasmid Plus Midi Kit (QIAGEN, Hilden, Germany).

### Large-scale yeast transformation

The large-scale yeast transformation was performed in three replicates, as described in Diss *et al.*^27^ (with slight modifications). To account for differences in library size, the WT ZF libraries and Ikaros-mimicry library were transformed using lower volumes than the DMS library. Exact volumes for each transformation are reported as pairs (e.g., WT ZF & Ikaros-mimicry *vs.* DMS), with values separated by a ‘/’ (e.g., 175/350 mL). BY4742 DDB1ΔB cells were streaked onto YPAD plates and incubated at 30°C for two days. For each replicate, 10 mL of liquid YPAD was inoculated with a single colony of the streaked-out cells and grown to saturation at 30°C and 200 rpm. Then, the OD_600_ was measured, and a culture of 175/350 mL YPAD was inoculated at an OD_600_ of 0.3 and grown for approximately 4.5 h at 30°C and 200 rpm to an OD_600_ of 1.2-1.6. Next, the cells were harvested by centrifugation for 5 min at 3,000 g, washed once with 50 mL ddH_2_O, and then washed with 50 mL SORB. Then the cells were resuspended in 7/14 mL SORB and incubated for 30 min at RT on a wheel. After incubation, 175/350 μL of 10 mg/mL preboiled salmon sperm DNA (Agilent Technologies, Santa Clara, CA) was added and mixed thoroughly, followed by the addition of 3.5/7.0 μg of the library to each replicate and thoroughly mixed. The three replicates of the DMS library were then split into two 7 mL portions, while the WT ZF library, with a lower volume, already contained 7 mL. 35 mL of Plate Mixture was added to each tube and incubated for 30 min at RT on a wheel. Next, 3.5 mL of DMSO (AppliChem, Darmstadt, Germany) was added, mixed thoroughly, and quickly placed in a water bath at 42°C for 20 min to undergo heat shock. The heat was homogenised within the tubes by inverting each tube five times at 1 min, 2 min 30 s, 5 min, 7 min 30 s, 10 min, 12 min 30 s, and 15 min. Next, the cells were centrifuged for 5 min at 1,700 g. The supernatant was removed by pouring, followed by a quick spin to collect any residual supernatant, which was then removed by aspiration. Then, the cell pellets were resuspended in 50 mL prewarmed Recovery medium and incubated at 30°C without agitation for 1 h. Next, the cells were harvested by centrifugation for 5 min at 3,000 g and then resuspended in 50 mL SC-Ura. The DMS library, which had been split into two, was then pooled again, centrifuged as before, followed by resuspension in 350/700 mL SC-Ura in a 2/5 L Erlenmeyer flask. Finally, 10 L of the culture was plated on SC-Ura plates to estimate transformation efficiency. The plates were incubated for 48 h at 30°C, and the cultures were grown at 30°C and 200 rpm for 48 h until saturation. The number of transformants varied between 1 and 5 million, ensuring a coverage of more than 10 transformants per variant in each replicate.

### GluePCA

The transformed cultures were grown to saturation in SC-Ura, followed by a second selection by inoculating 200/1,000 mL (WT ZF & Ikaros-mimicry library/DMS library, as described above) of SC-Ura/Ade/Met at an OD_600_ and the cultures were grown to an OD_600_ of 1.2 to 1.4. Then, the competition culture was inoculated at an OD_600_ of 0.05 in 200/1,000 mL SC-Ura/Ade/Met + 200 μg/mL MTX in 2% DMSO ± Compound. The competition culture was grown for 5 generations until an OD_600_ of 1.6 was reached. To prevent the accumulation of mutations and ensure reproducibility between the three replicates, the DMS library was grown for a maximum of 35 h, even if this was fewer than 5 generations. Input samples were harvested from the remaining cells of the second selection culture by centrifugation at 3,000 g for 5 min, followed by two washes with ddH_2_O and were stored at -20°C. The Output samples, which were harvested from the competition culture, were treated in the same way as the Input samples.

### DNA extraction

#### 1. Media

- Prime Buffer: 0.1 M EDTA-KOH pH 7.5, 10 mM DTT

- Zymolyase Buffer: 20 mM K-phosphate pH 7.2, 1.2 M Sorbitol, 0.4 mg/mL Zymolyase 20T (amsbio, USbiological), 100 μg/mL RNAse A

- Homemade P1: 50 mM Tris-HCl pH 8.0, 10 mM EDTA, 100 μg/mL RNAse A

- Homemade P2: 200 mM NaOH, 1% SDS

#### 2. Miniprep for 200 mL yeast cultures

Cell pellet was thawed at RT, resuspended in 4 mL Prime buffer and incubated at 30°C for 15 min. Then, the cells were centrifuged for 5 min at 2,500 g; the supernatant was discarded, and the pellet was resuspended in 4 mL of Zymolase buffer and was again incubated at 30°C for 1 h. Spheroplasting efficiency was evaluated under a microscope by mixing 4 μL of cells with 4 μL of 0.5% Triton X-100. When an efficiency of roughly 95% was reached, the cells were incubated further for 10 min and then harvested by centrifugation at 2,500 g for 5 min. Next, the cells were resuspended in 1.6 mL homemade P1 buffer, followed by adding 1.6 mL of homemade P2 buffer and mixed by inverting the tube several times and incubated for 10 min at RT. Then, 1.6 mL of QIAGEN N3 buffer (QIAGEN, Hilden, Germany) was added and mixed thoroughly. This was followed by centrifugation at maximum speed in an Eppendorf 5425 tabletop centrifuge for 20 min, and the supernatant was recovered. The entire supernatant was applied to a QIAGEN Miniprep column (QIAGEN, Hilden, Germany) on the QIAvac 24 Plus (QIAGEN, Hilden, Germany), washed once with 750 μL and twice with 200 μL QIAGEN wash buffer PE (QIAGEN, Hilden, Germany) and finally eluted in 50 μL QIAGEN elution buffer EB (QIAGEN, Hilden, Germany).

#### 3. Midiprep for 1 L yeast cultures

The beginning of the yeast Midiprep was the same as the yeast Miniprep, described above, with different buffer volumes up to the addition of homemade P2 buffer; the volumes were as follows: 20 mL Prime buffer, 20 mL Zymolyase buffer, 7.5 mL homemade P1 and 7.5 mL homemade P2 buffer. After incubation with homemade P2 buffer at RT for 10 min, 7.5 mL of pre-cooled QIAGEN P3 buffer (QIAGEN, Hilden, Germany) was added and the tube was inverted several times. Then, the sample was centrifuged at 4°C for 15 min at 3,000 g, and the supernatant was recovered into 50 mL centrifugation tubes (Nalgene, Rochester, NY, USA), followed by another centrifugation step at 4°C for 15 min at 15,000 g. The supernatant was recovered and filtered to remove remaining cell debris. Midiprep columns from the QIAGEN Midi Kit (QIAGEN, Hilden, Germany) were equilibrated with 10 mL QIAGEN QBT buffer (QIAGEN, Hilden, Germany). Then, the pre-filtered supernatant was added to the column, followed by two washes with 10 mL QIAGEN QC buffer (QIAGEN, Hilden, Germany). The plasmids were then eluted with 5 mL of QIAGEN QF buffer (QIAGEN, Hilden, Germany), and 3.5 mL of isopropanol was added. The mixture was mixed and centrifuged at 14,200 g for 15 min. The supernatant was discarded and 1 mL of freshly prepared 70% ethanol was added, then centrifuged again at 15,870 g for 1 min, the supernatant was discarded, and the pellet was left to dry at RT. Finally, the plasmids were resuspended in 150 μL of QIAGEN buffer EB (QIAGEN, Hilden, Germany).

#### 4. Plasmid concentration determination

To assess the molar plasmid concentration relative to genomic DNA, qPCR was performed on all samples using primers OGD241 (GCCTACATACCTCGCTCTGC) and OGD242 (CAACCCGGTAAGACACGACT), which bound to the plasmid backbone. The plasmid that served as the backbone during the cloning of the libraries (described above), was used to generate the standard curve. Concentrations for the standard curve were 0.4, 0.08, 0.016, 0.0032, 0.00064, 0.000128 and 0.0000256 ng/μL. The extracts were diluted 1:200, and qPCR was performed using the 2X SsoAdvanced Universal SYBR Green Supermix (Bio-Rad Laboratories, Hercules, CA) according to the manufacturer’s protocol.

### Sequencing library preparation

The plasmid did not contain the Illumina binding sites (SP1 and SP2); therefore, two PCRs were performed to introduce SP1 and 2. For the first PCR, primers ODL832 (CCCTACACGACGCTCTTCCGATCTTTCAGGAGCATCTGCTAGC) and ODL833 (TTCAGACGTGTGCTCTTCCGATCTAGCGTGACATAACTAATAAGC) were used. To ensure high coverage of input molecules, 280 million molecules of plasmid were used as input for the WT ZF library, while 1 billion molecules were used as input for the DMS library. The inserts were amplified for 16 cycles with 60°C annealing and 72°C extension for 15 s in 50 μL (0.3 mM dNTP mix, 0.15 μM primers, 0.5 U/reaction polymerase) reactions, followed by PCR clean-up with the QIAquick PCR Purification Kit (QIAGEN, Hilden, Germany) and eluted in 15 μL. 12 μL of the first reaction was used as input for the second PCR. To allow multiplexed sequencing, the NEBNext dual primer set 1/2 (New England Biolabs, Ipswich, MA) was used as primers for the second PCR. The second round of PCR was performed as the first round, but with a two-step 72°C annealing and extension for 30 s. For both PCRs, the KAPA HiFi Kits (Roche) were used. The samples were then further purified by 1% agarose gel electrophoresis, followed by gel extraction with the QIAquick Gel Extraction Kit (QIAGEN, Hilden, Germany), and eluted in 30 μL of QIAGEN EB buffer (QIAGEN, Hilden, Germany). Finally, the concentration was measured using the Qubit system (Invitrogen, Waltham, MA) and by qPCR with the KAPA Library Quantification Standards and Primers (Roche Sequencing Solutions Inc., Pleasanton, CA). All samples were pooled at equimolar ratios. The final pool was then further purified using AMPure XP beads (Beckman Coulter, Life Sciences, Brea, CA) according to the manufacturer’s protocol. The concentration was measured again using Qubit and qPCR. As a final quality control measure, the pool was loaded onto a Bioanalyzer (Agilent Technologies, Santa Clara, CA). For the WT ZF library, sequencing was performed on the Illumina HiSeq 2500 System (Illumina, San Diego, CA) using a 50 bp single-end sequencing run. The DMS and Ikaros-mimicry libraries were sequenced on the AVITI (Element Biosciences, San Diego, CA) with the 150 bp paired-end sequencing run. All runs were spiked with 25% PhiX phage library to increase sequencing quality.

### Sequencing data processing and binding score calculation

The fastq.gz files were read into R using the Mutscan^59^ package. Raw sequencing reads were filtered for an average Q score ≥ 30, and we silent mutations were allowed; all reads were then collapsed by amino acid sequence. The enrichment score (ES) was calculated for each variant I in replicate r as the ratio between its frequency before and after selection.

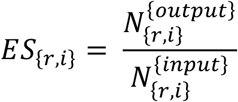

#### 1. WT ZF libraries & Ikaros-mimicry library

Sequences with <10 reads in one of the three DMSO replicates were excluded. After filtering, we retained 8,277 variants for the Single ZF library, 4,917 variants for the Dual ZF library and 1,919 variants for the Ikaros-mimicry library, which represent 99%, 93% and 96% of the respective libraries. Comparison of the replicates revealed high reproducibility, with Pearson correlations of approximately 0.85, 0.94 and 0.99 for the DMSO condition of the Single, Dual ZF and Ikaros-mimicry libraries, respectively. In these analyses, the ES was calculated with DMSO as input, and, to normalise, a limma trend (limma package^60^) was fitted to obtain the final log_2_FC. In addition, the DMSO condition was also used to correct for abundance, since little to no growth was expected without the compound.

#### 2. DMS library

Sequences with <10 reads in one of the three Input replicates were excluded. After filtering, we retained 76,730 variants, which accounted for 85% of the whole library. The reproducibility of the Input for the DMS library was also good, with a Pearson correlation of approximately 0.75. The ES score was calculated as described above with Mutscan using the limma method. To correct for abundance, a symmetric LOESS regression model (span = 0.8) was fitted with the compound log₂FC as a function of the DMSO control. The resulting residuals were extracted and interpreted as the abundance-corrected binding.

### Cellular ZF degradation assay

Degradation assays were performed as previously described^61^. Commercially available 293T cells (Thermo Fisher Scientific, Waltham, MA) were cultured in Dulbecco’s modified Eagle’s medium supplemented with 10% FBS and 1% penicillin–streptomycin. The ZF constructs were synthesised and cloned into a lentiviral expression plasmid (Artichoke, Addgene #73320) as fusions to GFP with an IRES-separated mCherry as an expression control. To produce the lentivirus, 550,000 Lenti-X 293T cells (Takeda, 632180) were seeded in a six-well plate with 3 mL of growth medium. The following day, a plasmid solution containing 500 ng pMD2.G (Addgene #12259), 1000 ng psPAX2 (Addgene #12260), and 1000 ng ZF-Artichoke plasmid was prepared in Opti-MEM (Thermo Fisher Scientific, Waltham, MA, 31985070), which resulted in a total volume of 37.5 μL. A second solution consisting of 9 μL Opti-MEM and 3.75 μL TransIT-LT1 (Sigma, St. Louis, MO) was prepared, incubated at room temperature for 5 min, and then gently mixed into the plasmid solution. The transfection mix was incubated at room temperature for 30 min before being added to the Lenti-X cells. The cells were allowed to grow for 48 h, after which the virus-containing supernatant was centrifuged (800 g, 2 min), aliquoted in 300 μL volumes, and stored at -80°C.

For transduction, 3 million 293T cells were resuspended in 2.7 mL of growth medium in a six-well plate, and 300 μL of virus was added. The plate was centrifuged at 800 g for 1 h at 37°C, then incubated for 48 h. Cells were selected with puromycin (1 μg/mL) for a week. For the flow cytometry experiments, 25,000 cells were seeded in a 96-well plate format in 0.09 mL of growth medium and incubated for 16 h. Cells were then treated with compounds (diluted in 10 μL growth medium) at the indicated concentration and duration. The medium was aspirated, and the cells were washed once with PBS. Then, 0.05% trypsin was added. After 5 min of incubation, cells were collected via centrifugation (400 g for 2 min), resuspended in 100 μL PBS and measured using a BD LSR II. Data were analysed with FlowJo (v10.4.0, BD Biosciences) to determine the mean fluorescence signal ratio GFP-to-mCherry for each well. Gating strategy can be found in **Extended Fig. 8a**. Treatment ratios were then normalised to the corresponding DMSO control to assess degradation of each ZF construct.

### Protein Expression

#### 1. Expression and Purification of CRBN and DDB1ΔB

Human FL CRBN (residues 1-442) and human DDB1ΔB (residues 1-395/706-1140 with GNGNSG as a linker in between) were subcloned into pAC-derived vectors^62^. For CRBN containing an N-terminal Strep II tag and DDB1ΔB with a 6xHis tag, the Bac-to-Bac system (Thermo Fisher Scientific, Waltham, MA) was used to co-express them in 4 L cultures of Trichoplusia ni High Five (Hi5 cells). Cells were cultured at 27°C for two days after infection and collected by centrifugation. The cell pellet was resuspended in lysis buffer (50 mM Tris-HCl pH 8.0, 500 mM NaCl, 10% glycerol, 0.5 mM TCEP, 0.1% Triton X-100, 5 mM MgCl_2,_ 0.5 mM CaCl_2_, 1x protease inhibitor cocktail (Sigma) and DNase) and lysed by sonication. After ultracentrifugation, the supernatant was collected, and proteins were purified by Strep-Tactin affinity chromatography (IBA), followed by ion-exchange chromatography (Cytiva) and further purified by size-exclusion chromatography (SEC; Superdex 200; Cytiva) in SEC buffer (50 mM HEPES pH 7.4, 200 mM NaCl, 10% Glycerol and 0.5 mM TCEP). The purified proteins were concentrated and stored at -80°C.

#### 2. Expression and Purification of ZF Proteins

Human Ikaros^ZF23^ (residues 141-243(Δ197-238)), SALL4^ZF123^ (residues 361-591), and Helios^ZF123^ (residues 95-195) were cloned into pAC-derived vectors with an N-terminal Strep II tag and expressed with the Bac-to-Bac system in 4 L Hi5 cell cultures. Cells were cultured for two days at 27°C, harvested by centrifugation and resuspended in lysis buffer (50 mM HEPES pH 7.4, 700 mM NaCl, 10% glycerol, 0.5 mM TCEP, 0.1% Triton X-100, 5 mM MgCl_2_, 0.5 mM CaCl_2_, 1x protease inhibitor cocktail (Sigma) and DNase), and purified by Strep-Tactin affinity chromatography in SEC buffer (50 mM HEPES pH 7.4, 500 mM NaCl, 0.5 mM TCEP, 10% glycerol). The proteins were further purified by SEC (Superdex 200), concentrated, and finally stored at -80°C.

### Cryo-EM sample preparation

CRBN-DDB1ΔB was mixed with 100 μM pomalidomide or ALV2 (MedChemExpress) and a 4-fold molar excess of the ZF Proteins in 150 μL of binding buffer (25 mM HEPES pH 7.4, 0.5 mM TCEP and 150 mM NaCl). The mixed proteins were then cross-linked using GraFix^63^. Here, the protein complexes were layered onto a 10-30% (v/v) glycerol gradient (25 mM HEPES pH 7.4, 150 mM NaCl, 0.5 mM TCEP and 100 μM pomalidomide/ALV2) with an increasing concentration of glutaraldehyde (0-0.25%; EMS) and subjected to ultracentrifugation (Beckman SW40Ti rotor, 30 krpm, 16 h at 4°C). The gradient was harvested in 100 μL fractions and analysed by SDS-PAGE; the peak fractions were then pooled. Then, the glycerol was removed with Zeba Desalting Columns (Thermo Fisher Scientific, Waltham, MA) and the sample was concentrated to 2-4 μM. 4 μL of sample was applied to Quantifoil holey carbon grids (R1.2/1.3 200-mesh, Quantifoil Micro Tools), which were glow-discharged with a Solarus plasma cleaner (Gatan) for 120s in a H_2_/O_2_ environment. After sample application, the grids were blotted for 4 s at 4°C and 100% humidity using a Vitrobot Mark IV (FEI) and immediately plunged into liquid ethane.

### Cryo-EM data collection (see Table S1 Statistics for details)

All datasets, except for the SALL4^ZF123^∼CRBN∼DDB1^ΔB^ dataset, which was recorded with a Glacios (Thermo Fisher Scientific, Waltham, MA) operating at 200 kV, were collected on a Cs-corrected (CEOS GmbH) Titan Krios (Thermo Fisher Scientific, Waltham, MA) operating at 300 kV.

### Cryo-EM image processing

Real-time evaluation, and acquisition with EPU 3.0 (Thermo Fisher Scientific, Waltham, MA), was performed with CryoFlare v1.10^64^. Drift was corrected with RELION 3 motion-correction implementation^65^, where a motion-corrected sum of all frames was generated with and without applying a dose-weighting scheme. All datasets were further processed in cryoSPARC v4^66^ as indicated in each Extended Data figure, including two-dimensional (2D) and 3D classification, 3D refinement, particle polishing, and CTF refinement. Reported resolution values for all reconstructions are based on the gold-standard Fourier shell correlation curve (FSC) at the 0.143 criterion^66^, and all related FSC curves are corrected for soft-mask effects, using high-resolution noise substitution^67^. DeepEMhancer was used for sharpening^68^.

### Model building and refinement

For modelling Helios^ZF123^, we used PDB 9E2U^20^ as a template. For Ikaros^ZF23^, we extracted the coordinates from the AlphaFold3^46^ prediction for Ikaros and combined them with PDB 4CI3^2^, removing the WD-repeat β-propeller B. The two models were fitted into the cryo-EM map using ChimeraX^69^ (fit-in-map tool). Finally, the models were refined using ISOLDE^70^ in combination with adaptive distance restraints and Phenix^71^. Side chains were corrected in Coot and ISOLDE if necessary.

### Human proteome surface fingerprint database

The publicly released AF2 proteome database contains 23,391 human proteins^72^. We split each AF2 model into individual domains based on a cutoff of 15 Å in their corresponding predicted aligned error (PAE) matrices. In addition, the low-confidence disordered regions (i.e., pLDDT lower than 70) at both N- and C-termini of each domain were removed. PAE matrices are not available for models already split into overlapping fragments due to the length limitation (1,400 amino acids) in the AF2 proteome database; they were thus processed using the Merizo algorithm^73^ into individual domains with the pLDDT filter set to 50. This ultimately led to a database of 75,684 domains of the human proteome. MaSIF fingerprints were then computed for all the structural domains as previously described^48^. Briefly, the molecular surfaces of these human proteome proteins are decomposed into overlapping radial patches with a 12 Å radius, capturing, on average, nearly 400 Å^2^ of surface area. The chemical and geometric features of these surface patches are then passed through a neural network to generate the surface descriptors.

### MaSIF-mimicry search

MaSIF-mimicry search leverages the established MaSIF-seed search pipeline, as described previously^74^, with modifications made to perform alignments based on similarity rather than complementarity to the target patch. Specifically, we used MaSIF-seed descriptors without the inversion of the input features and normal of the target patch before performing alignment, which ultimately enables the search for similar patches rather than complementary. To score the similarity between patches, the descriptor distance between each vertex of the query patch and its nearest neighbour in the target patch in the post-alignment state was computed. The descriptor distance score (DDS) aggregates all the descriptor distances according to the following formula, as previously described^75^:

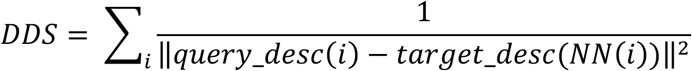

where *DDS* is the descriptor distance score, *i* indexes interacting patches of the first protein, and *NN(i)* returns the index of the spatially nearest neighbour on the other protein. Higher scores mean higher similarity. *DDS* was then normalised to a score in the range from 0 to 1 using the following equation:

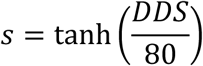

where *s* is referred to as the MaSIF similarity score, representing the similarity level between the two protein surface patches.

### Benchmarking Single ZF dataset

For benchmarking the Single ZF dataset, the 5 patches closest to the glycine residue in the G-loop motif of ZNF692 ZF4 (PDB ID: 6H0G) were selected. These were compared against all surface patches of ZF domains in the MaSIF human proteome database. The ability of interface mimicry to predict binding was evaluated against other methods, including full-length sequence similarity, interface sequence similarity, and embedding similarity from a protein language model. Full-length sequence similarity was measured by performing a global pairwise alignment with PairwiseAligner from Biopython, using the BLOSUM62 matrix. The similarity scores for all pairs were normalised by dividing by the maximum value, resulting in a range of 0 to 1. The interface pseudo-sequence consists of 5 key residues at positions corresponding to the interface residues in ZNF692 ZF4 (Q418, E420, G423, F424, and T425) that contact CRBN-pomalidomide. We used residues that are at the same relative positions to the two cysteines and two histidines in the Single ZF sequence as the interface pseudo-sequence (i.e., the residue before the first cysteine, the residue after the first cysteine, and the first 3 residues after the second cysteine). These pseudo-sequences were then encoded into numerical vectors using the BLOSUM62 scheme to capture evolutionary relationships of amino acids, and we used the cosine similarity between the encoded sequences to measure the interface sequence similarity. In addition, each construct’s full-length sequence in the Single ZF dataset was encoded using ESM-2^76^ (650M) to obtain its sequence representation (i.e., embedding), and again, the similarity was measured with the cosine similarity between encoded sequences.

### Proteome-wide search

For the proteome-wide search, the query patches were limited to the nearest 5 patches to the glycine residues in the G-loop motif of Ikaros (PDB ID: 6H0F), GSPT1 (PDB ID: 6XK9), and Ck1α (PDB ID: 5FQD). Given the large size of the MaSIF human proteome database, during the search process, each protein surface in the database was uniformly down-sampled by selecting every third patch (i.e., a 1 to 3 down-sampling size on the surface). After performing the alignment between the query and target patch, the same transformation matrix was applied to the entire query protein structure to calculate the number of Cα and heavy atom clashes between the query proteins and CRBN. The poses with a MaSIF similarity score higher than the cut-off of 0.4 were retained, and the pose with the highest MaSIF similarity score for each domain hit was selected as the representative prediction for downstream analysis.

### Synthetic DNA library design

MaSIF mimicry search of Ikaros (PDB ID: 6H0F) yielded 110,356 unique post-alignment states with a MaSIF similarity score > 0.4 and TM-score < 0.5, corresponding to 25,947 structural-domain models predicted to bind CRBN-pomalidomide in an Ikaros-like binding mode. To experimentally screen these predictions by GluePCA, we designed a synthetic DNA library encoding the top candidates. We limited the library to 2,000 genes of ≤300 bp (≤86 residues), where for each unique domain model, we selected the post-alignment state with the highest MaSIF similarity score for further analysis.

Each selected binder model was first aligned to the Ikaros ZF2 based on interface similarity. Interface residues on each binder were defined within 4.5 Å of the CRBN-pomalidomide complex. Domains longer than 86 residues were truncated: an overlapping 86-residue windows that collectively cover all interface residues are first generated. Outlier residues were removed iteratively based on the z-score of their sequence index to maintain the 86-residue limit (Extended Data Fig. 7d). Each truncated fragment was scored for predicted thermodynamic stability by EvoEF2^77^. The sub-domain with the most favourable ΔG was advanced for structural filtering. After further interface and quality assessments (see **Analysis of truncated models and filtering criteria**), we retained 2,020 truncated models. The protein sequences of the top 1,959 truncated models, ranked by MaSIF similarity score, were encoded into DNA sequences using the same method as mentioned above (see Library curation section).

### Analysis of truncated models and filtering criteria

The following metrics were computed on each truncated model in complex with CRBN-pomalidomide and used to filter out low-quality candidates:

- Normalised EvoEF2^77^ score: the EvoEF2-predicted folding ΔG (kJ/mol) of the truncated model divided by its residue count. Models with normalised score < – 3 kJ/mol/residue were accepted, removing variants likely to be unstable or unfolded.
- Interface residues lost due to truncation: the number of interface residues in the full-length model (within 4.5 Å of CRBN-pomalidomide) absent from the truncated model’s interface. Models with ≤ 2 residue losses were accepted, ensuring that critical contacts were kept.
- Normalised SASA: the total solvent-accessible surface area (Å²) of the truncated binder divided by its residue count. Models with an area of ≤ 122 Å² per residue were accepted to exclude overly exposed or elongated structures, which are indicative of poor folding.
- Mean distance of interface residues to termini: the average sequence distance (in residues) from each interface residue to the nearest N- or C-terminus. Models with a mean distance of 15 residues or more were accepted, removing interfaces that were too close to termini and prone to be problematic due to fusion of the reporter domains in the GluePCA screen.
- Number of binder interface residues: count of residues within 4.5 Å of CRBN–pomalidomide. Models with four or more interface residues were accepted to ensure sufficient contacts.
- Percentage of intracellular interface residues: based on DeepTMHMM^78^ topology predictions for the full-length sequence; models with ≥ 80% of interface residues predicted intracellular were accepted, excluding transmembrane or extracellular binders.
- Ligand-binder interface area: the buried SASA of pomalidomide upon binding the truncated model (isolated SASA minus complexed SASA). Models with a buried area of ≥ 120 Å² were accepted to ensure direct ligand interaction.
- Mean interface pLDDT score: average pLDDT of interface residues. Models with a score of ≥ 79.2 were accepted to exclude low-confidence interfaces.
- Based on the secondary structure assigned by STRIDE^79^, models with ≥ 4 contiguous segments (helix, strand, or loop) and with ≥ 20% structured residues were accepted to remove overly simplistic or unstructured folds.
- Normalised geodesic length: the longest 3D geodesic path (Å) divided by the cube root of the model’s voxel-computed volume (Å³), with accepted values ≤ 7,495 to exclude excessively elongated or artefactual structures.

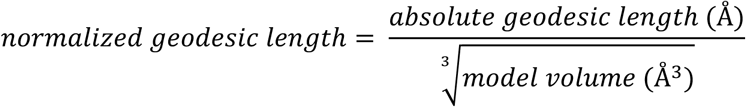

### Structural and interface space analysis

To examine the structural diversity of the 50 surface mimicry hits from the bonsai library, we performed all-against-all TM alignment. We used the TM score to represent the differences in the backbone structures. Additionally, we combined the structures of CRBN and pomalidomide with these hits to generate ternary complexes and performed an all-against-all comparison to assess interface diversity. The predicted interfaces should be highly similar to each other, as they were all retrieved based on interface mimicry to either Ikaros ZF2 or ZNF629 ZF4. Therefore, we used an orthogonal tool, iAlign^80^, to calculate the IS-score, which considers the geometric distance and the conservation of interfacial contact patterns between each pair of ternary complexes to measure the interface similarity.

### Druggability analysis

To assess the druggability of each hit, we employed the druggability score and Prank score predicted by Fpocket software for protein domains within the human proteome from a previous study^54^. In the original study, unstructured regions were filtered out, and the druggability was analysed only for protein domains within autonomous structural units defined in Pfam. For domains in our results that lack a druggability or Prank score, we assigned a score of 0 in the table.

### Figure preparation

Structural figures were prepared with UCSF ChimeraX (v.1.3). Degradation assay plots were prepared with GraphPad Prism (v.10.4.2), schematics were done in Adobe Illustrator 2025, molecules were drawn in Chemdraw and the surface similarity plot was done in Python (3.13.7) with seaborn. All other graphs were prepared with R^81^, specifically with the package ggplot2^82^.

---

**Extended Data Figure 1:**
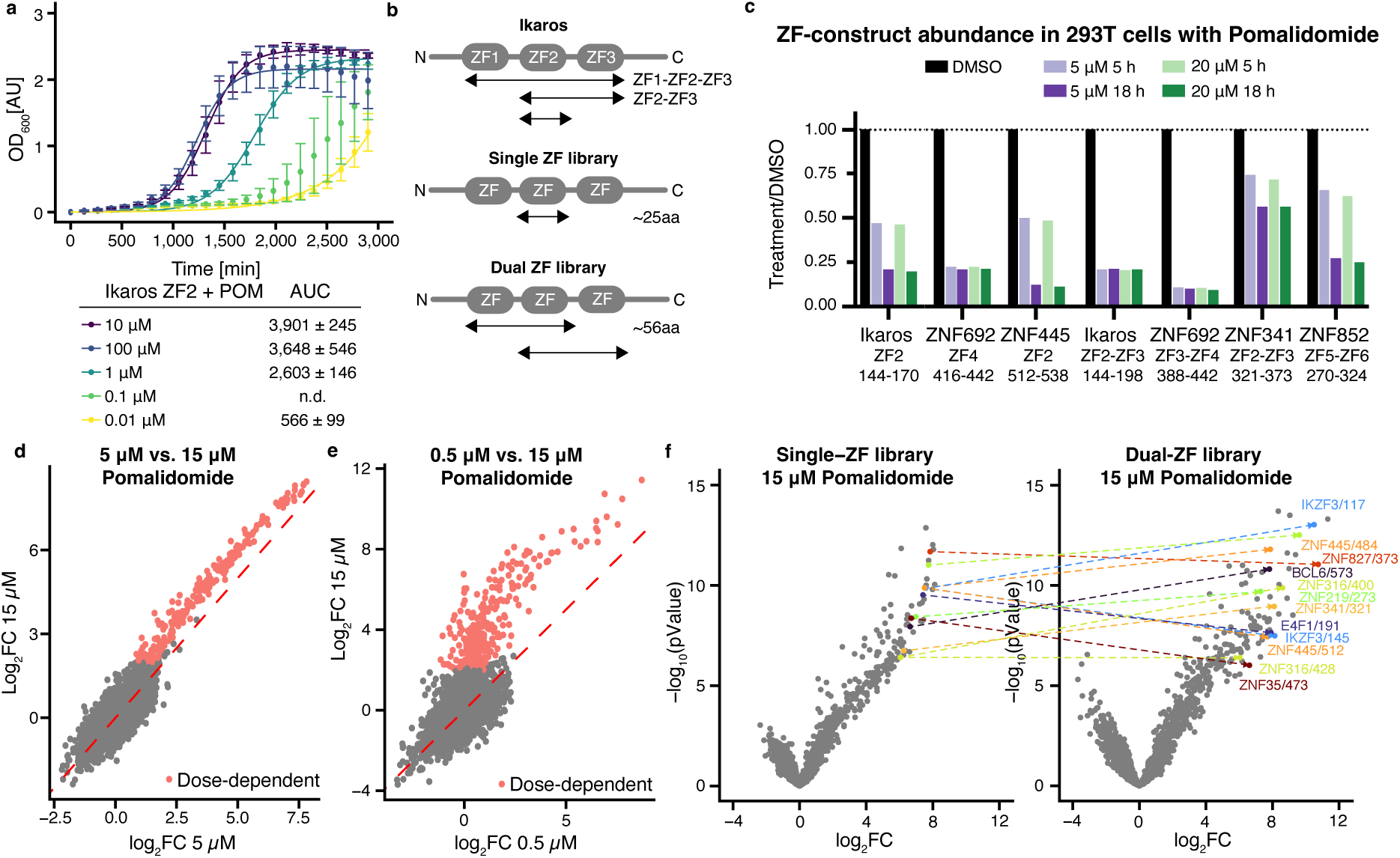
Compound-dependency to identify true binders. **(a)** Single-clone GluePCA in yeast with Ikaros ZF2 at different pomalidomide concentrations. (Biological replicates = 3, each point indicates the mean of the experimental replicates, error bars indicate the SD) **(b)** Schematic of tested Ikaros ZF constructs and the Single/Dual ZF library. **(c)** 293T cells expressing the indicated ZF-GFP-IRES-mCherry construct were treated for 5 and 18 hours with DMSO or pomalidomide at the indicated doses. Cells were then analysed by flow cytometry to quantify the ratio of GFP over mCherry fluorescence, and treatments were normalised to DMSO. **(d)** Log_2_FC of Single ZF library at 5 µM pomalidomide plotted against 15 µM pomalidomide. The red line indicates the diagonal (x = y). The dose-dependent binders are indicated as red dots. **(e)** Log_2_FC of Dual ZF library at 0.5 µM pomalidomide plotted against 15 µM pomalidomide. The red line indicates the diagonal. The dose-dependent binders are indicated as red dots. **(f)** Comparison of the single ZF and dual ZF GluePCA screen. Connected points are single-ZF constructs (left) with their corresponding dual ZF construct or constructs (right).

**Extended Data Figure 2:**
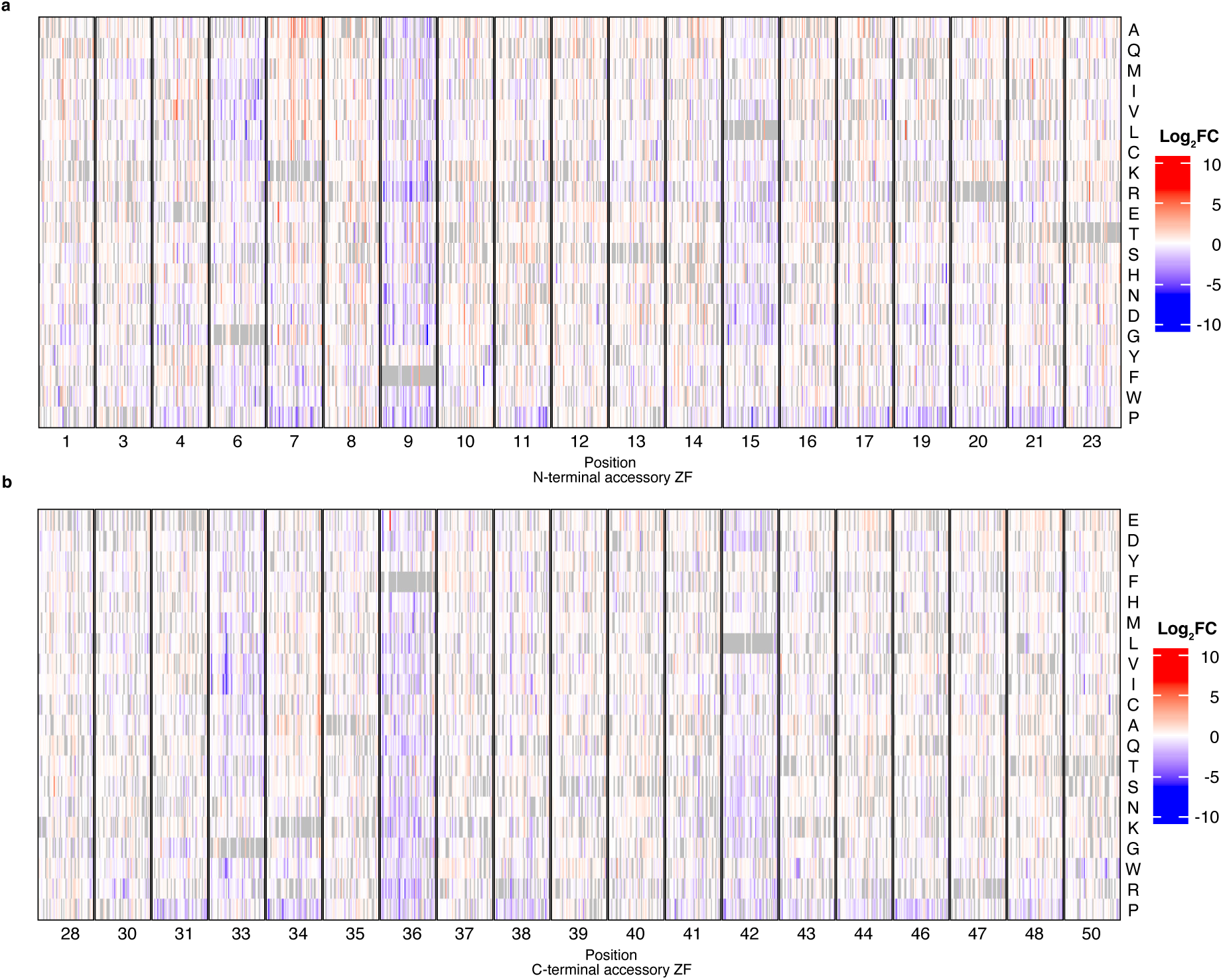
Measured enrichment of all N- and C-terminal accessory ZFs. (a) Heatmap of all aligned N-terminal aZF. The black rectangle indicates a single position; each column represents a ZF construct at the corresponding position. Each row indicates the mutant amino acid. (Biological replicates = 3, each rectangle indicates the mean of the experimental replicates) (b) Heatmap of all aligned C-terminal aZF. The black rectangle indicates a single position; each column represents a ZF construct at the corresponding position. Each row indicates the mutant amino acid (biological replicates = 3, each rectangle indicates the mean of the experimental replicates).

**Extended Data Figure 3:**
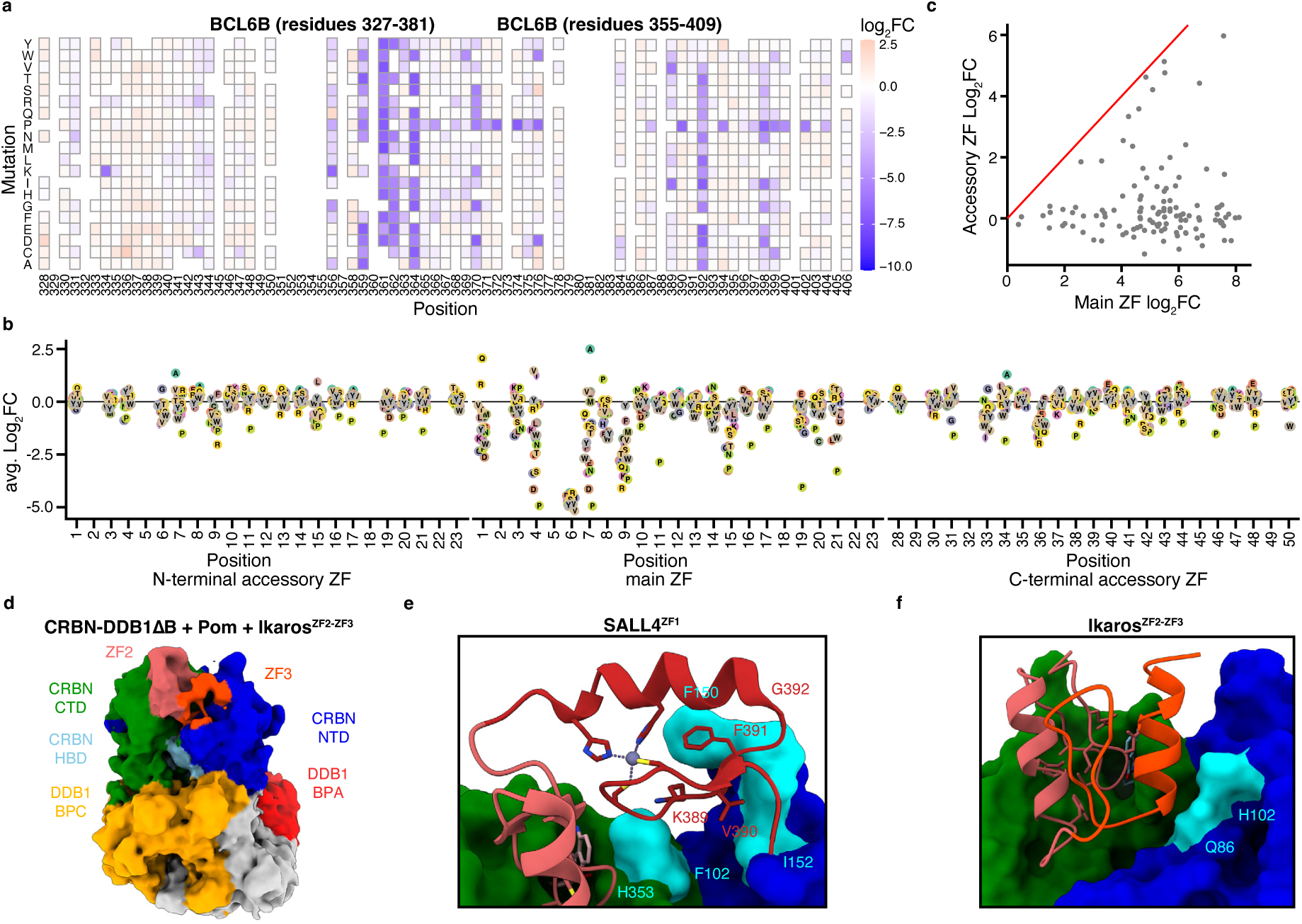
N- and C-terminal accessory ZF facilitates ZF∼CRBN binding through favourable PPI. (a) Example heatmaps of the DMS of overlapping constructs BCL6B (residues 327-381) and BCL6B (residues 355-409) in GluePCA. (b) Average log_2_FC of all mutations at each position after alignment of the mZF (c) Log_2_FC of the mZF of each ZF used in the DMS screen plotted against its aZF. (d) Cryo-EM structure of Ikaros^ZF23^ bound to CRBN-DDB1ΔB-pomalidomide. (e) SALL4^ZF12^ bound to CRBN-pomalidomide complex model from Ma *et al.*^33^. (f) Tentative Ikaros^ZF23^ model. ZF2 in red and ZF3 in orange. In cyan are the proposed CRBN interaction residues.

**Extended Data Figure 4:**
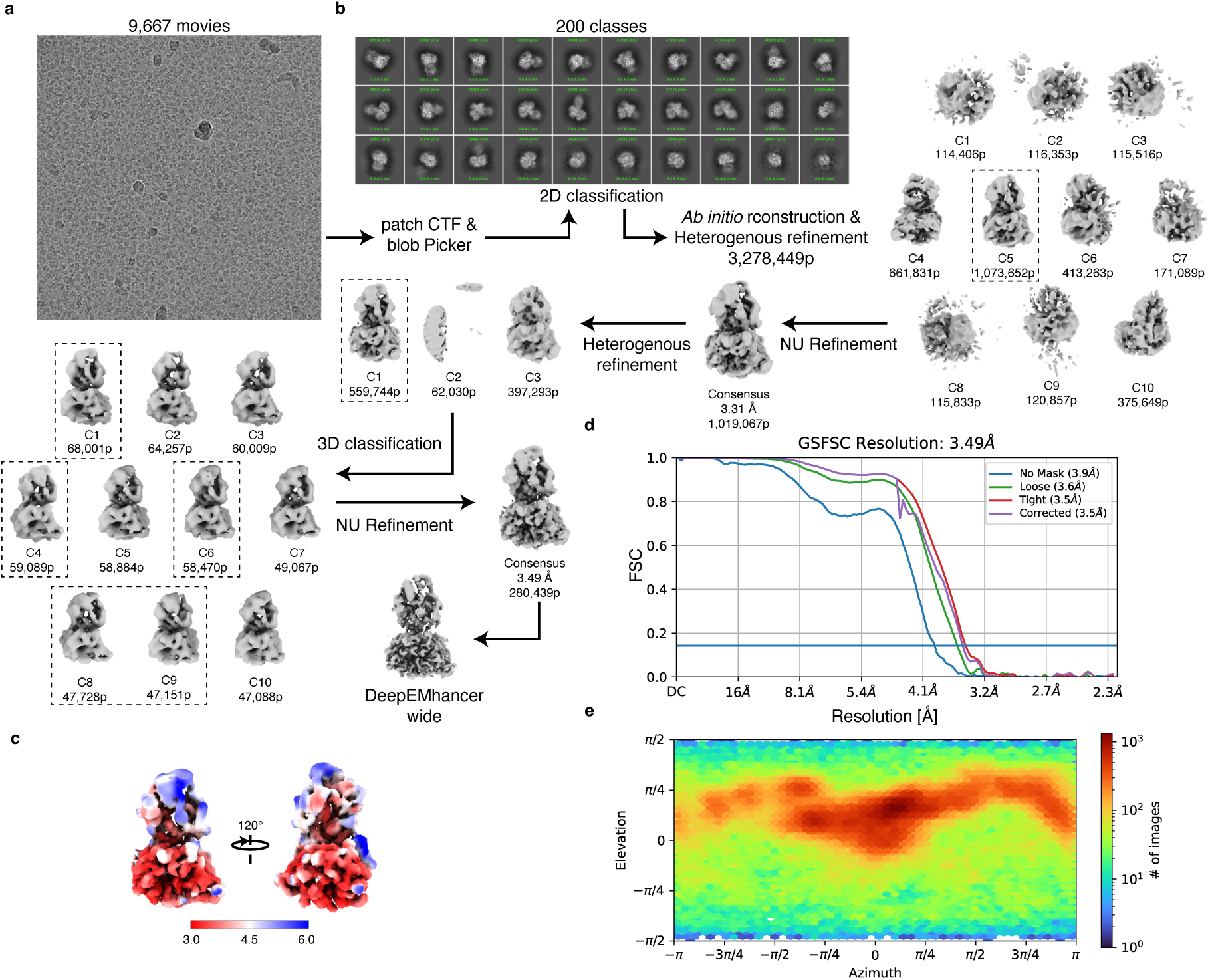
Processing of CRBN-DDB1ΔB pomalidomide SALL4^ZF12Helix^ cryo-EM structure. (a) Representative cryo-EM micrographs for the dataset. (b) The whole analysis was done in CryoSparc. Particles were picked using the blob picker, followed by an *ab initio* reconstruction and Heterogenous refinement (the black dashed line indicates further processed volumes). The particles were further processed by 3D classification and finally sharpened with DeepEMhancer. (c) Calculated local resolution of the final map. (d) Gold-standard FSC curve for the 3.49 Å resolution map. (e) Angular distribution for the particles leading to the 3.49 Å resolution map.

**Extended Data Figure 5:**
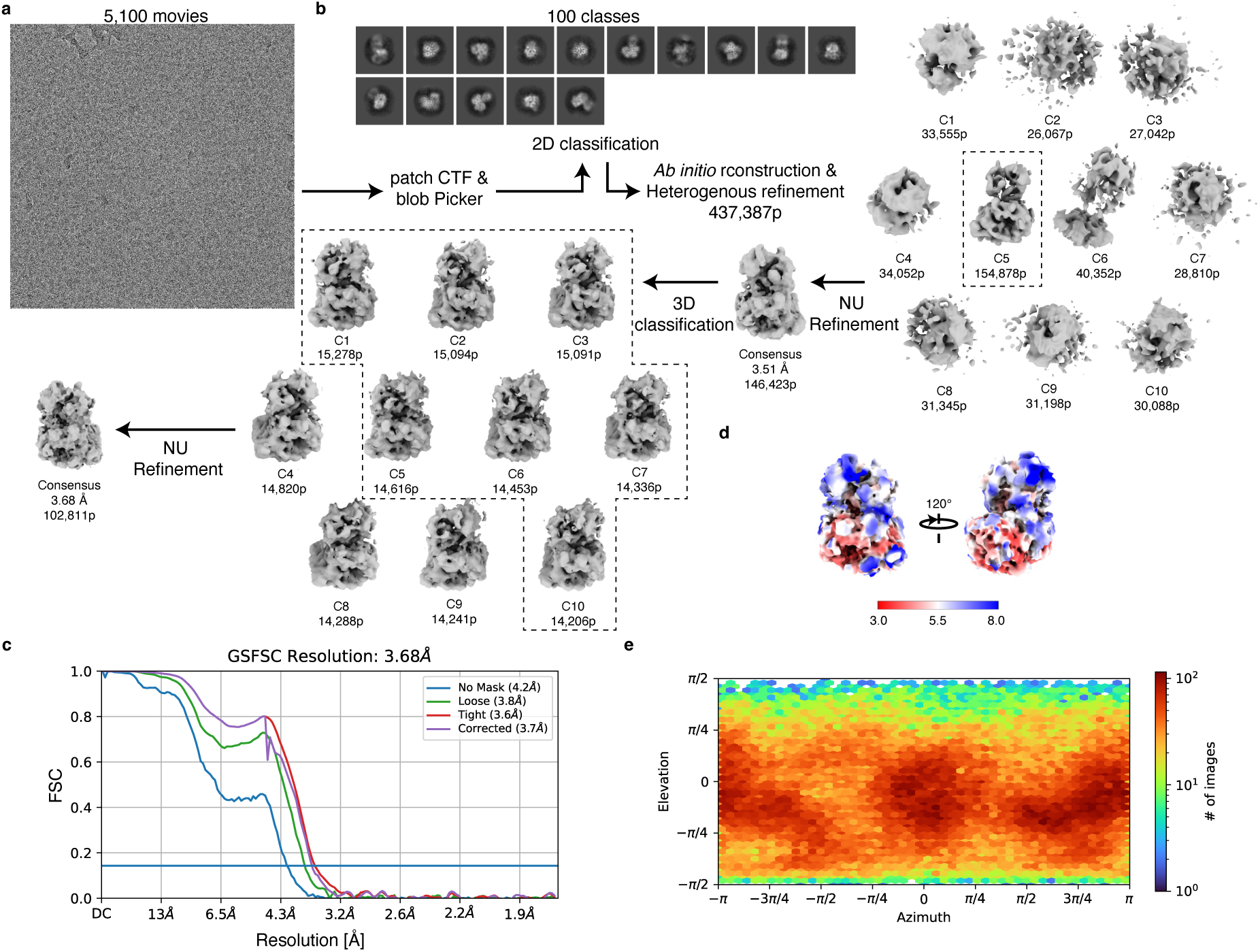
Processing of CRBN-DDB1ΔB pomalidomide Ikaros^ZF23^ cryo-EM structure. (a) Representative cryo-EM micrographs for the dataset. (b) The whole analysis was done in CryoSparc. Particles were picked using the blob picker followed by an ab initio reconstruction and Heterogenous refinement (the black dashed line indicates further processed volumes). The particles were further processed by 3D classification and finally refined with a Non-Uniform (NU) refinement (c) Gold-standard FSC curve for the 3.68 Å resolution map. (d) Calculated local resolution of the final map. (e) Angular distribution for the particles leading to the 3.68 Å resolution map.

**Extended Data Figure 6:**
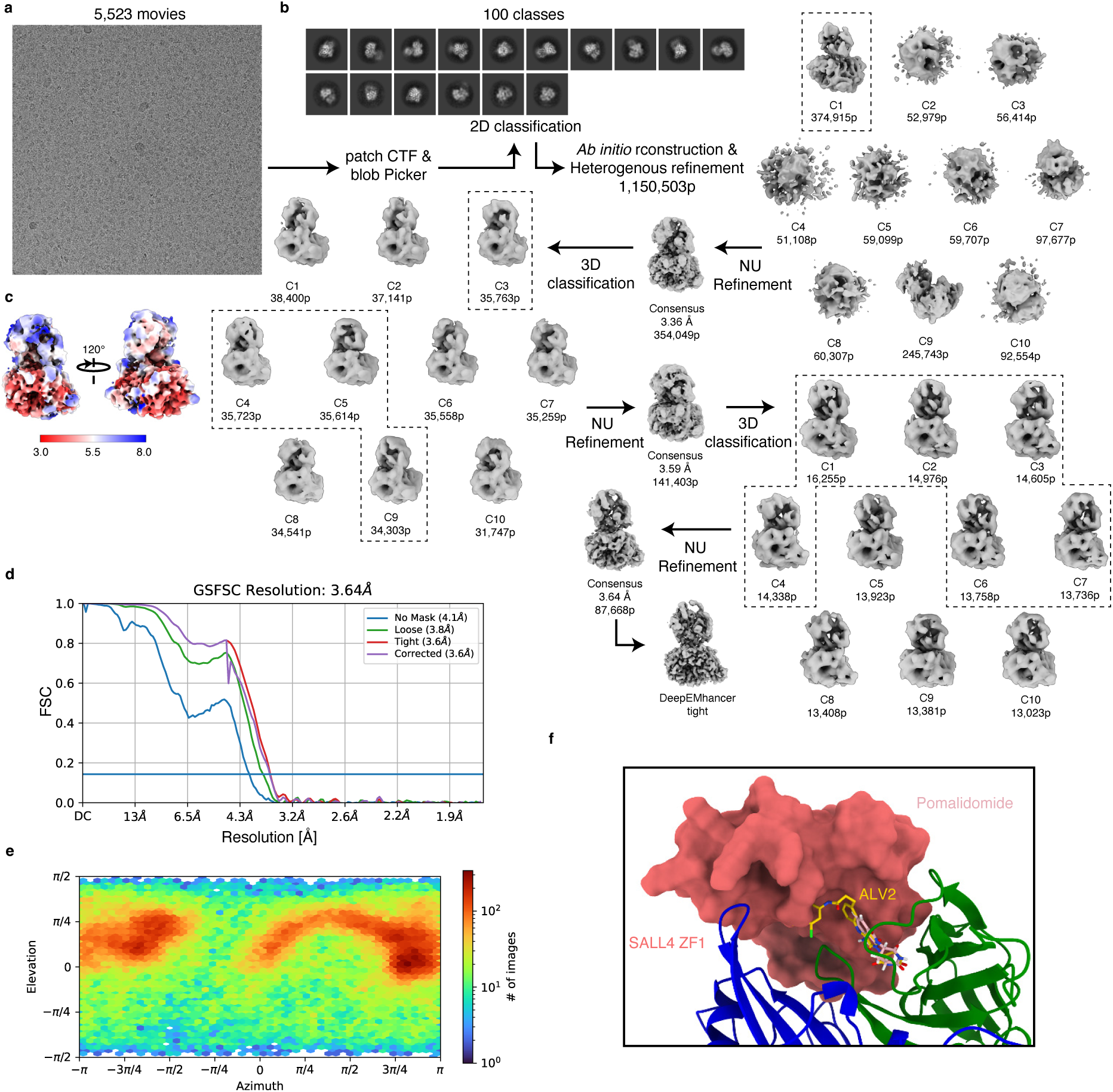
Processing of CRBN-DDB1ΔB pomalidomide Helios^ZF123^ cryo-EM structure. (a) Representative cryo-EM micrographs for the dataset. (b) The whole analysis was done in CryoSparc. Particles were picked using the blob picker, followed by an ab initio reconstruction and Heterogenous refinement (the black dashed line indicates further processed volumes). The particles were further processed by 3D classification and finally sharpened with DeepEMhancer. (c) Calculated local resolution of the final map. (d) Gold-standard FSC curve for the 3.64 Å resolution map. (e) Angular distribution for the particles leading to the 3.64 Å resolution map. (f) Comparison of SALL4 ZF1 and ALV2 bound to CRBN, which is predicted to clash.

**Extended Data Figure 7:**
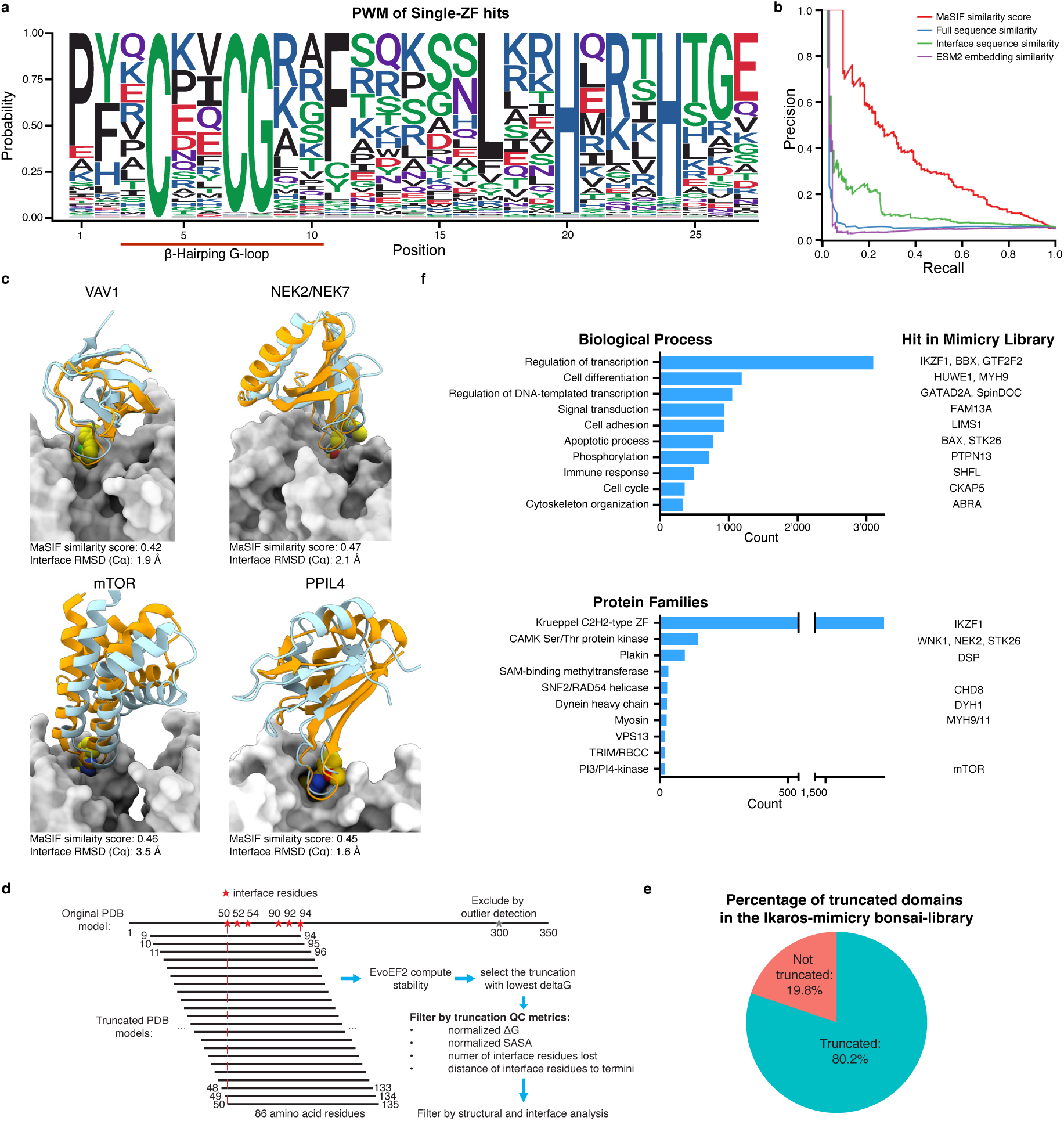
Surface mimicry can predict novel targets from a diverse set of biological processes and protein families. **(a)** Position weight matrix (PWM) of all Single-ZF GluePCA hits. **(b)** Precision-recall curves comparing various homology-based prediction methods on the Single ZF dataset benchmark, using ZNF692 ZF4 as the reference. **(c)** Comparison of crystal structures (VAV1, NEK7, mTOR and PPIL4 (orange)) to the binding modes predicted (cyan) by MaSIF-mimicry search, which is performed based on interface mimicry to their respective query structures (GSPT1, Ikaros, and ZNF692). The crystal structures (PDB: 9NFR, 9H59, 9NGT and 9DWV) are first aligned to the CRBN in their respective query structures (PDB: 6XK9, 6H0F, and 6H0G). The interface RMSD measures the average distance of interface Cα atoms between the predicted structure and the crystal structure. **(d)** Library design strategy: predicted binders exceeding 86 amino acid residues were truncated by generating overlapping 86-residue windows that collectively cover all interface residues and selecting the truncation window with the most favourable predicted folding ΔG. The truncated candidates were then filtered by various metrics ensure the structural integrity, interface quality, and biological sensibility of the candidates. **(e)** Ikaros-mimicry Bonsai-library composition. In red the percentage of non-truncated, FL domains. In blue the percentage of truncated domains. **(f)** Top 10 most represented biological processes and protein families in MaSIF-mimicry.

**Extended Data Figure 8:**
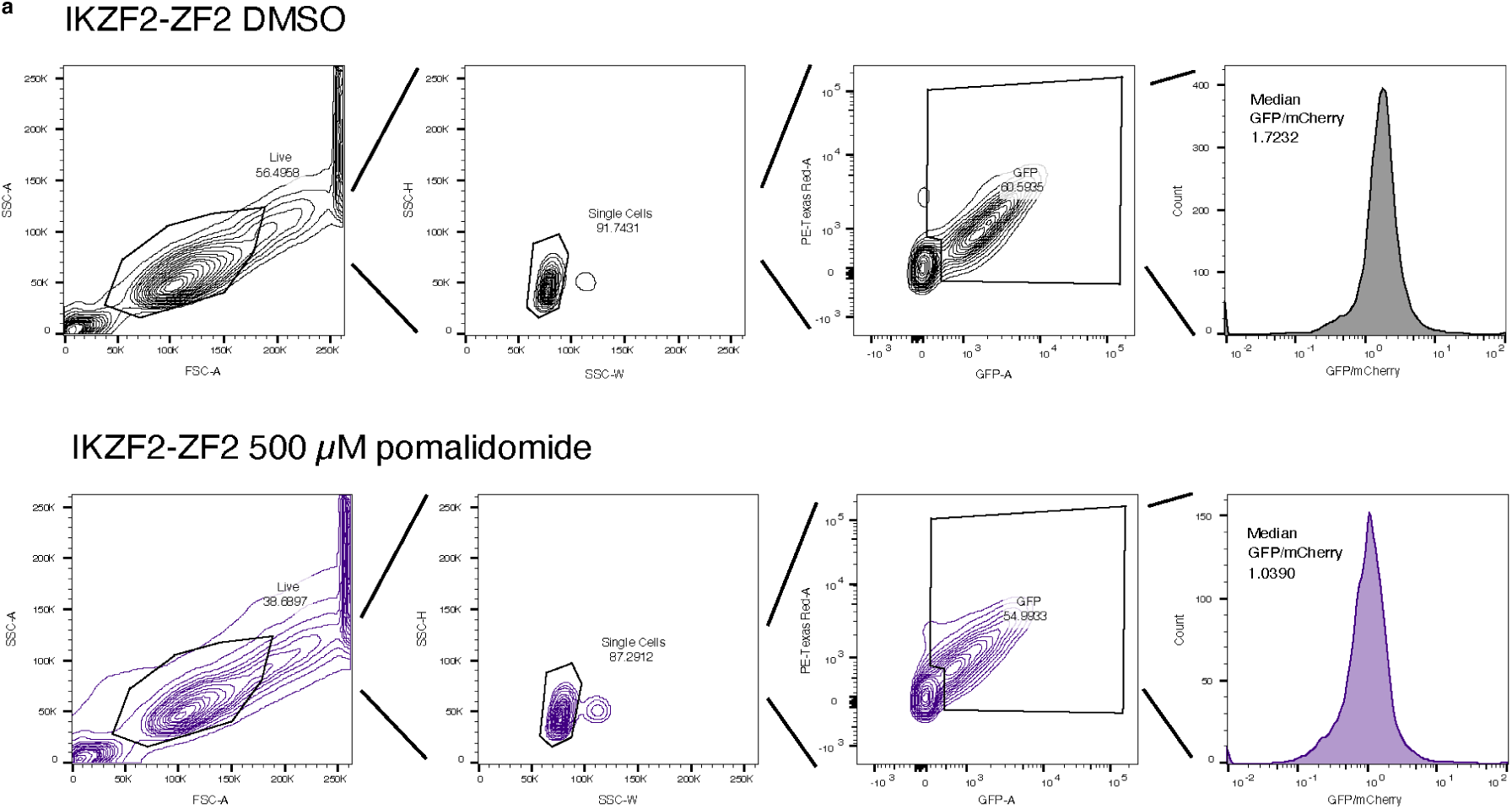
Schematic of the gating strategy used in flow cytometry. (a) Illustration of the gating strategy in flow cytometry

**Table S1:** Cryo-EM data collection, refinement, and validation statistics. 1) Sharpening was done with DeepEMhancer

|  | **SALL4**<br>**ZF12Helix** | **Ikaros ZF23** | **Helios ZF123** |
| --- | --- | --- | --- |
| **Data collection and processing** |  |  |  |
| EMDB ID | EMD-55106 | EMD-55107 | EMD-55108 |
| PDB ID | 9SQ4 | 9SQ5 | 9SQ6 |
| Detector | Falcon 4 | Falcon 4i | Falcon 4i |
| Magnification | 120,000 | 75,000 | 75,000 |
| Voltage (kV) | 200 | 300 | 300 |
| Electron exposure (e<sup>-</sup>/Å) | 50 | 50 | 50 |
| Defocus range (μm) | -2.0 to -0.8 | -2.0 to -0.8 | -2.0 to -0.8 |
| Pixel size (Å) | 0.84 | 0.845 | 0.845 |
| Symmetry imposed | C1 | C1 | C1 |
| Initial particle images (no.) | 3,278,449 | 437,387 | 1,150,503 |
| Final particle images (no.) | 280,439 | 102,811 | 87,668 |
| Map resolution (Å) | 3.49 | 3.68 | 3.64 |
| FSC threshold | 0.143 | 0.143 | 0.143 |
| Map resolution range (Å) | 3-9 | 3-15 | 3-12 |
| **Refinement** |  |  |  |
| Initial models used (PDB codes) |  |  |  |
| Model resolution (Å) | 3.49 | 3.68 | 3.64 |
| FSC threshold | 0.143 | 0.143 | 0.143 |
| Map sharpening B factor (Å) | N.A.<sup>1</sup> | N.A.<sup>1</sup> | N.A.<sup>1</sup> |
| Model composition |  |  |  |
| Non-hydrogen atoms | 9992 | 10037 | 9907 |
| Protein residues | 1256 | 1263 | 1241 |
| Ligands | LIG: 1<br>ZN: 2 | LIG: 1<br>ZN: 3 | LIG:1<br>ZN: 4 |
| B factors (Å) |  |  |  |
| Protein | 121.27 | 202.71 | 81.55 |
| Ligands | 138.58 | 294.41 | 122.43 |
| R.m.s deviations |  |  |  |
| Bond lengths (Å) | 0.004 | 0.003 | 0.004 |
| Bond angles (°) | 0.662 | 0.567 | 0.695 |
| Validation |  |  |  |
| MolProbity score | 0.93 | 1.37 | 1.22 |
| Clashscore | 0.68 | 2.64 | 1.8 |
| Poor rotamers (%) | 0.19 | 0.48 | 0.39 |
| Ramachandran plot |  |  |  |
| Favored (%) | 96.67 | 95.33 | 95.95 |
| Allowed (%) | 3.25 | 4.50 | 3.88 |
| Disallowed (%) | 0.08 | 0.17 | 0.17 |
| **Model-to-data fit** |  |  |  |
| CCmask | 0.75 | 0.79 | 0.68 |
| CCbox | 0.76 | 0.89 | 0.67 |
| CCpeaks | 0.72 | 0.71 | 0.65 |
| CCvolume | 0.75 | 0.79 | 0.69 |

